# A pair of dopaminergic neurons DAN-c1 mediate *Drosophila* larval aversive olfactory learning through D2-like receptors

**DOI:** 10.1101/2024.01.15.575767

**Authors:** Cheng Qi, Cheng Qian, Emma Steijvers, Robert A. Colvin, Daewoo Lee

## Abstract

The intricate relationship between the dopaminergic system and olfactory associative learning in *Drosophila* has been an intense scientific inquiry. Leveraging the formidable genetic tools, we conducted a screening of 57 dopaminergic drivers, leading to the discovery of DAN-c1 driver, uniquely targeting a pair of dopaminergic neurons (DAN) in the larval brain. While the involvement of excitatory D1-like receptors is well-established, the role of D2-like receptors (D2Rs) remains underexplored. Our investigation reveals the expression of D2Rs in both DANs and the mushroom body (MB) of third instar larval brains. Silencing D2Rs in DAN-c1 via microRNA disrupts aversive learning, further supported by optogenetic activation of DAN-c1 during training, affirming the inhibitory role of D2R autoreceptor. Intriguingly, D2R knockdown in the MB impairs both appetitive and aversive learning. These findings elucidate the distinct contributions of D2Rs in diverse brain structures, providing novel insights into the molecular mechanisms governing associative learning in *Drosophila* larvae.

## Introduction

Learning defines a behavioral change that results from acquiring information about the environment, and memory refers to the process by which the information is encoded, stored, and later retrieved. Learning and memory forms the basis for higher brain functions, including cognition and decision making, which shapes our individuality^1^. On the cellular and physiological level, learning and memory is achieved by neuroplastic changes in circuits, including neuronal excitability and synaptic plasticity. Usually distinct types of neurotransmitters, such as dopamine, modulate these changes.

Dopamine (DA) plays an important role in many mammalian brain functions, including motor functions, motivation, reinforcement, addiction, and learning and memory^2–4^. Dopaminergic neurons (DAN) are mainly located in the mesencephalon: DANs in the substantia nigra (SN) are responsible for motor functions, while those in the ventral tegmental area (VTA) are important in reward, addiction, learning and memory^3,5^. Dopamine achieves its functions via two families of G protein-coupled receptors (GPCR): excitatory D1-like and inhibitory D2-like receptors^4^. All D1-like receptors are located post-synaptically; in contrast, D2-like receptors both function post-synaptically and play an important presynaptic role, regulating dopamine release through negative feedback^4^. All these receptors are important in mammalian associative learning^6,7^. D1-like receptors elevate intracellular cAMP by activating adenylyl cyclase (AC) via Gα_s_, while D2-like receptors repress cAMP by inhibiting AC via Gα_i/o_. cAMP activates protein kinase A (PKA), leading to the phosphorylation of DARPP-32 (dopamine and cyclic AMP-regulated phosphoprotein, 32kDa), ion channels, and CREB (cAMP response element-binding protein). In addition, dopamine receptors also activate the PLC-PKC, MAPK, and CaMKII pathways^2–4,8–10^.

Although mammalian studies reveal mechanisms more relevant to human beings, the complexity of the nervous system impedes our understanding about the basic or more universal principles of learning and memory applicable generally to all nervous systems^11^. With a simple central nervous system (CNS) and powerful genetic tools, the fruit fly *Drosophila melanogaster* has become a popular model organism in learning and memory research^12,13^. With conserved genes in dopamine metabolism and signaling^14^, as well as fundamental similarities in the olfactory circuitry compared to mammals^15,16^, *Drosophila* can perform olfactory associative learning in both larvae^17–19^ and adults^20–24^. Olfactory associative learning is a type of classical conditioning in which flies are trained under positive or negative reinforcement paired with an odorant. Different from the naïve reaction to the odorant, flies approach the odorant after being trained with rewards (e.g., sucrose; appetitive)^21^, but avoid the odorant when trained with punishments (e.g., electric shock, bitter taste chemicals; aversive)^20^. Several genes related to the cAMP-PKA signaling pathway, including *dunce* (*dnc*)^25^ and *rutabaga* (*rut*)^21^, are expressed in the mushroom body (a center for *Drosophila* learning and memory) in both larval and adult brains^26,27^. Mutations of these genes lead to learning deficiencies, indicating their roles in larval^17,18^ and adult olfactory learning^21,25^.

*Drosophila* larvae offer several advantages for studying olfactory learning compared to adults. Notably, compared to neural circuits underlying olfactory learning, their simpler neural circuitry^28^, characterized by fewer olfactory receptor neurons (ORNs)^29^, projection neurons (PNs)^30^, mushroom body neurons (MBNs)^31^, and dopaminergic neurons (DANs)^32^, facilitates the elucidation of underlying mechanisms. Additionally, larvae exhibit simpler behavioral patterns, facilitating experimental manipulations and observations. Furthermore, their translucent cuticles enable convenient application of techniques such as optogenetics^33^, further enhancing the experimental versatility of larval studies.

Like in mammalian brains, dopamine achieves its functions via four dopamine receptors in flies, two D1-like receptors dDA1^34^ (or Dop1R1) and DAMB^35^ (or Dop1R2), one D2-like receptor D2R^36^ (or Dop2R), and one non-canonical receptor DopEcR^14,37^. dDA1 is mainly found in the mushroom body^38,39^ and is necessary for appetitive and aversive olfactory learning in larvae^39^ and adults^40^. D2R in the adult mushroom body is necessary for anesthesia-resistant memory^41^. In addition, D2R in GABAergic anterior paired lateral (APL) neurons is known to secure aversive conditioning in adult flies^42^. Although D2R expression has been reported in the ventral nerve cord^43^, neither its expression in larval brains, nor its functions in larval olfactory learning have been investigated.

By using a GFP-tagged D2R strain, we detected the expression pattern of D2R in the third-instar larval brain, specifically in dopaminergic and mushroom body neurons. Knockdown of D2Rs in a pair of DANs, DAN-c1, impaired aversive learning, while knockdown of D2R in mushroom body neurons led to deficits in both aversive and appetitive learning. These results revealed that D2Rs in distinct brain structures mediate different learning tasks. The newly discovered roles of D2R in the larval brain provides new insights into the mechanisms underlying larval associative learning.

## Results

### Distinct dopaminergic neurons innervate different compartments of the mushroom body

The connectome of larval learning circuitries has been investigated in both first- and third-instar larvae^28,44^. The mushroom body serves as a primary learning center in *Drosophila*, which is composed by αβ, α’β’, and γ neurons in adult brains^45^. In larvae, axons from γ neurons bifurcate and form the vertical and medial lobes, as αβ and α’β’ neurons are not mature^31,46,47^. These lobes are divided into 11 compartments (refer to Figure 1) based on the coverage of neurites from both mushroom body extrinsic neurons (MBEN) and mushroom body output neurons (MBON)^28^. Around 21 dopaminergic neurons are found in each third-instar brain hemisphere, and categorized into 4 clusters: DM1 (dorsomedial), pPAM (primary protocerebral anterior medial, or DM2)^48^, DL1 (dorsolateral), and DL2^32^. DL1 neurons project to the vertical lobe^39^, while pPAM neurons innervate the medial lobe^48^ (refer to Figure 1a-c).

**Figure 1.**
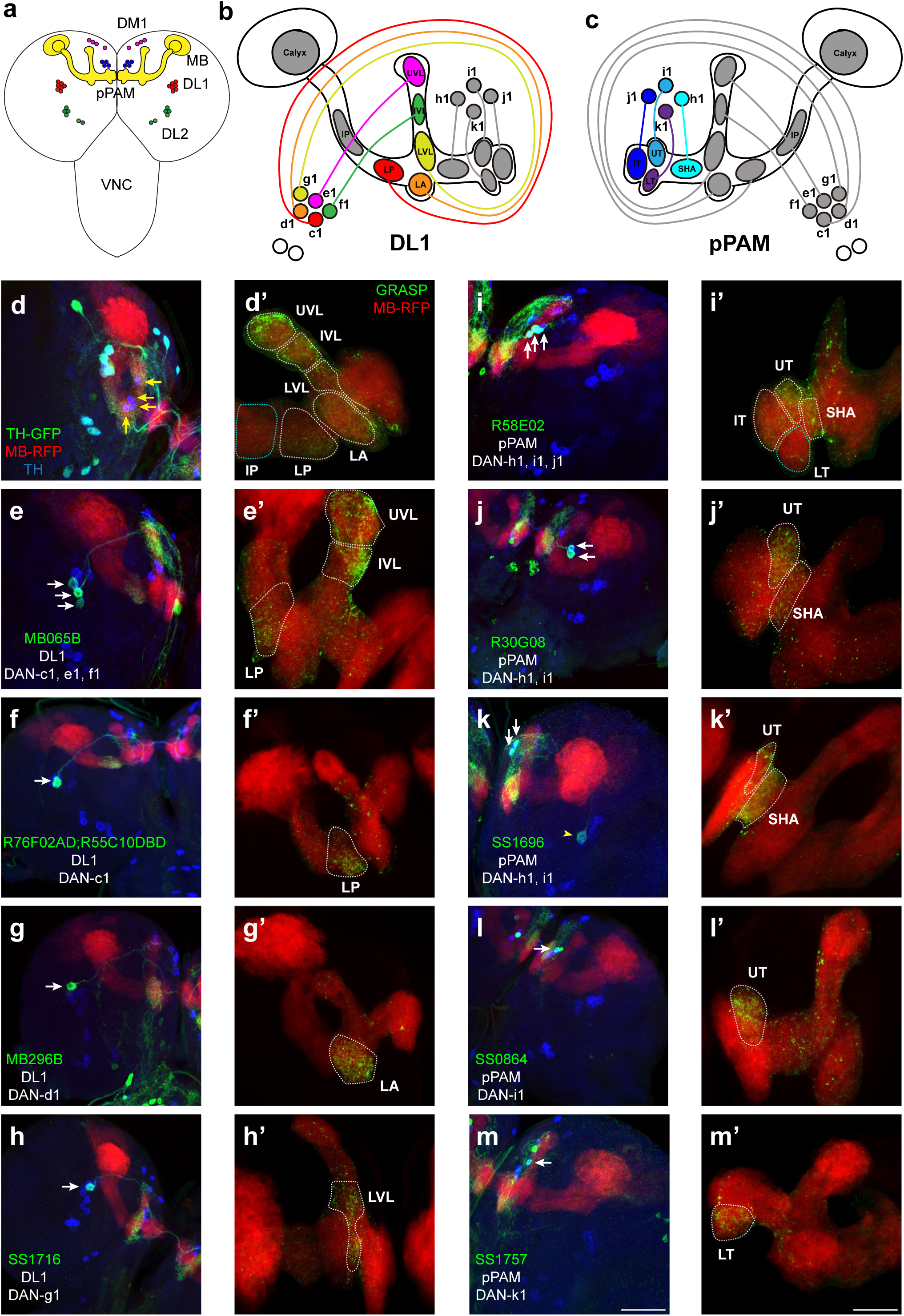
Identification of driver strains for a pair of dopaminergic neurons in the *Drosophila* larval brain. **(a)** A schematic diagram shows the dopaminergic neuron (DAN) clusters and the mushroom body (MB) in the third-instar larval brain. **(b)** & **(c)** Schematic diagrams show the innervation patterns from distinct DANs to different compartments in the MB. All 11 MB compartments are shown, note that there is no synapse formed from dopaminergic neurons (DANs) to the calyx and intermediate peduncle (IP). DANs in DL1 (b) and in pPAM (c). **(d-m)** Drivers covering DANs in the DL1 cluster (d-h). Drivers covering DANs in the pPAM cluster (i-m). The first column shows drivers covering distinct dopaminergic neurons. Neurons under the drivers are labeled by GFP, the mushroom body (MB) is labeled by RFP, and DANs are marked by tyrosine hydroxylase (TH) antibody. The second column (d’-m’) shows the GRASP signals from DANs under the driver to the corresponding compartments in the MB. The green channel represents the GRASP signals, and RFP marks the morphology of the MB. In the first column, white arrows mark the DANs under the driver strains. Yellow arrows in (d) show the pPAM neurons not labelled by TH-GAL4 driver strain. The yellow arrowhead in (k) showed the DL1 neuron not innervating the MB. Scale bars: 50 µm for the first column, and 20 µm for the second column. ***Abbreviations:*** DL, dorsolateral; DM, dorsomedial; IP, intermediate peduncle; IT intermediate toe; IVL, intermediate vertical lobe; LA, lateral appendix; LP, lower peduncle; LT, lower toe; LVL, lower vertical lobe; pPAM, primary protocerebral anterior medial; SHA, shaft; UT, upper toe; UVL, upper vertical lobe. *(Note) GFP expression patterns in the entire larval CNS by GAL4 driver strains used in this study can be found in Figure S1. N numbers for each strain can be found in Table S2*.

In this study, we wanted to functionally identify individual DANs that mediate larval olfactory learning. The first step was to search for DAN-specific driver strains that mark a few dopaminergic neurons, which subsequently can be used to target genetic manipulations of corresponding neurons. A total of 57 driver strains identifying dopaminergic neurons were screened in this study (Table 1). These strains were chosen based on previous studies, either identifying a pair of dopaminergic neurons in larvae, or identifying only several in adult brains and indicating the potential of identifying a pair of DANs in larvae. TH-GAL4, a traditional dopaminergic neuronal driver^49^, identifies all dopaminergic neurons except those in pPAM (Figure 1d, Figure S1a). Split-<u>G</u>FP <u>r</u>econstitution <u>a</u>cross <u>s</u>ynaptic <u>p</u>artners (GRASP) technique was used to investigate the “direct” synaptic connections from DANs to the mushroom body (MB), in which portions of GFP were specifically expressed in corresponding neurons (Figure S2d)^50^. GRASP results showed neurons under TH-GAL4 formed synapses in the vertical lobe and lower peduncle (white dash lines in Figure 1d’), consistent with previous electron microscopy data^44^. We found three DAN driver strains that identify a pair of dopaminergic neurons in the third-instar larval brain. DAN driver R76F02-AD;R55C10-DBD identifies a pair of dopaminergic neurons innervating the lower peduncle (LP), which would be DAN-c1 based on previous published nomenclature^28^ (Figure 1f). MB296B driver identifies the dopaminergic neurons (DAN-d1) projecting to the lateral appendix (LA) (Figure 1g), as well as many non-dopaminergic neurons. SS1716 driver identifies a pair of dopaminergic neurons forming synapses in the lower vertical lobe (LVL), indicating it is DAN-g1 (Figure 1h).

**Table 1.**
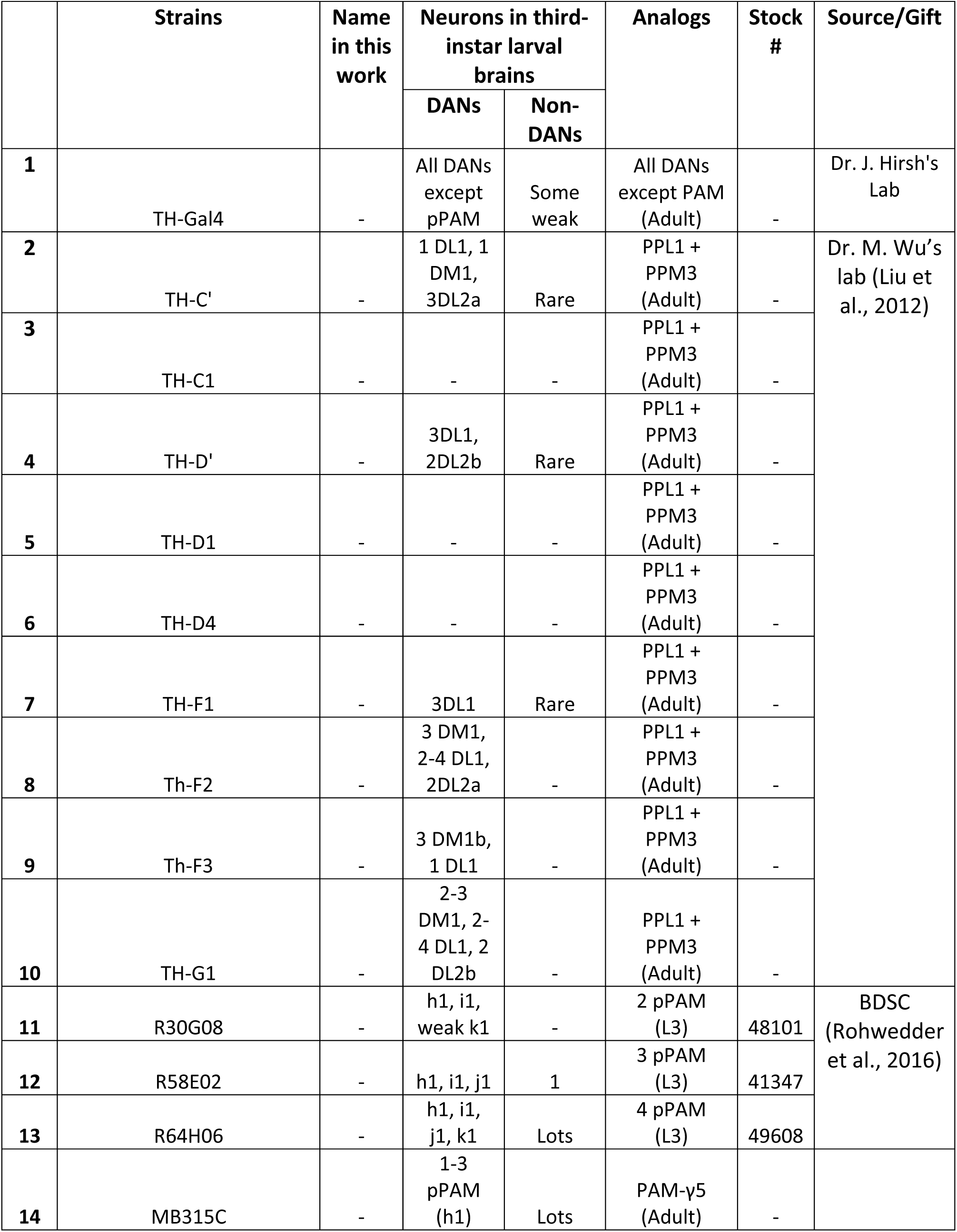
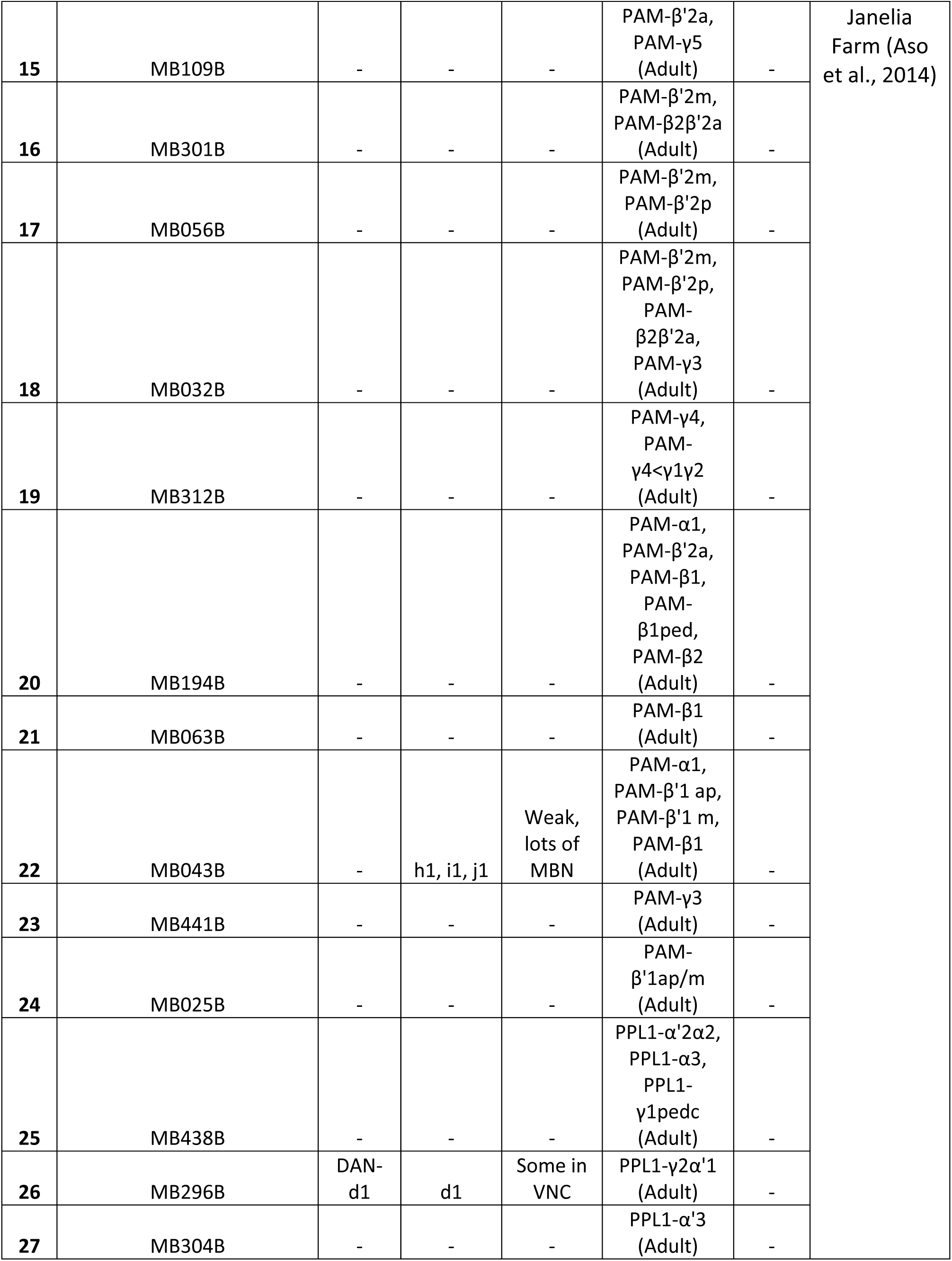
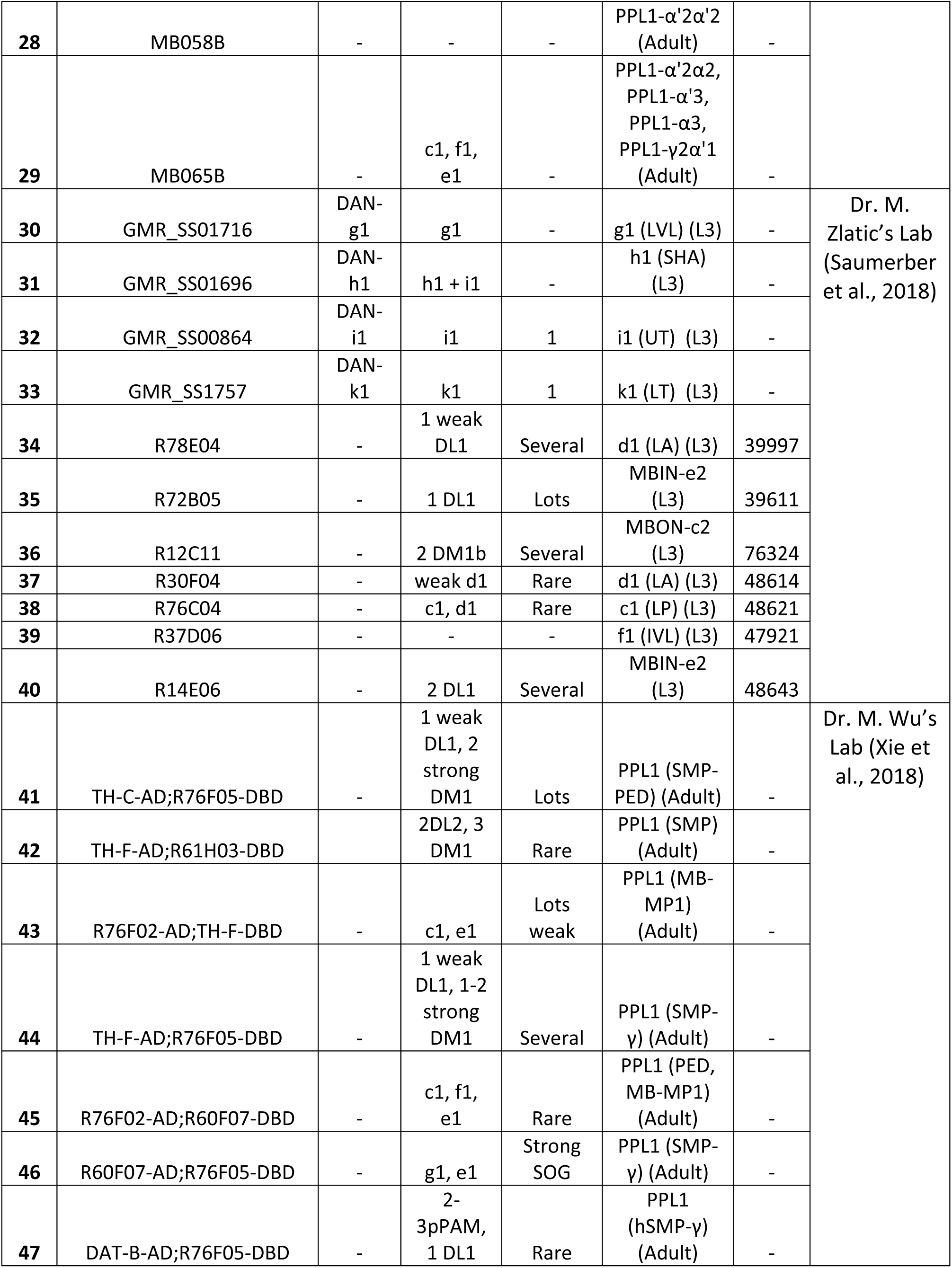
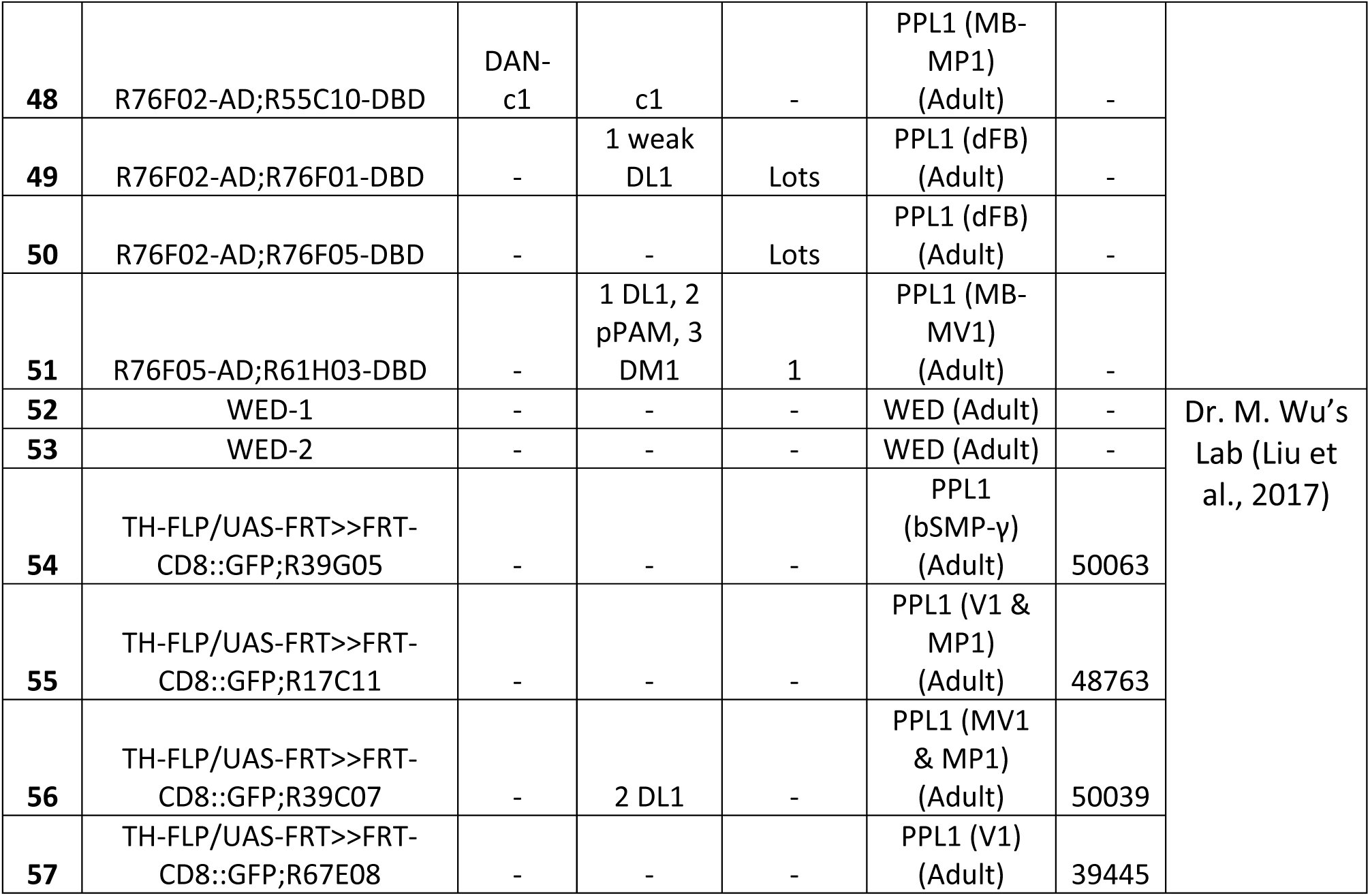
Driver strains screened for dopaminergic neurons in the third-instar larval brain. Strains are listed with their published names and names used in this work. The numbers of dopaminergic and non-dopaminergic neurons in the third-instar larval brain are described. The identities of dopaminergic neurons from distinct clusters are also listed, especially for those in DL1 and pPAM. The analogs column lists the labelled neurons in previous publications. Source/Gift column shows the original papers in which these strains were described, as well as the laboratories these strains were obtained from. Several, 2-5 neurons; some, 6-10 neurons; lots, >10 neurons

In pPAM, R58E02 driver identifies DAN-h1, i1, and j1, innervating the shaft (SHA), intermediate toe (IT) and upper toe (UT) (Figure 1i), and R30G08 driver identifies DAN-h1 and i1 (Figure 1j). As described in a previous report^28^, SS864 driver identifies DAN-i1, innervating the upper toe (Figure 1l), and SS1757 driver identifies DAN-k1 which innervates the lower toe (LT) (Figure 1m). In contrast, SS1696 driver identifies not only DAN-h1, but also i1 and one DL1 neuron not innervating the mushroom body (Figure 1k).

In summary, our results show that five DL1 and four pPAM DANs innervate nine distinct mushroom body compartments in a one-to-one pattern (Figure 1b and c). DL1 neurons innervate the vertical lobe and peduncle, while pPAM neurons project to the medial lobe. The neuronal driver strains screened can be used to investigate the roles of individually identified DANs in larval olfactory learning.

### Dopamine release from DAN-c1 mediates larval aversive learning

Dopamine plays an important role during olfactory associative learning in both adults^51,52^ and larvae^17,18^. In adults, dopaminergic neurons in PPL1 regulate aversive learning^53–56^, while those in PAM mediate reward signals in appetitive learning^57–59^. In larvae, DL1 neurons innervating the vertical lobe and the peduncle are required for aversive learning^18,39^, while those in pPAM projecting to the medial lobe are involved in appetitive learning^48^.

In Figure 1 and Table 1, three driver strains identifying distinct pairs of dopaminergic neurons in DL1 were discovered, which could be candidates to investigate their roles in larval aversive learning. The R76F02-AD;R55C10-DBD strain identifies MB-MP1 in the adult brain^60^, which is a dopaminergic neuron involved in adult aversive learning^61^. Based on the analysis with 22 brain samples, we observed this driver strain labels one neuron per hemisphere in the third-instar larval brain (Figure 2a-d, Figure S1c, Table S3). Using a UAS-DenMark;UAS-sytGFP strain, its dendrites were labeled with RFP and axonal terminals were marked by GFP. Its dendrites were localized in the dorsomedial protocerebrum (dml), and its axonal terminals located in the lower peduncle of the mushroom body (Figure 2a and c), with GRASP results supporting the existence of synapses in this compartment (Figure 1f’). All these characteristics are consistent with the previously published nomenclature^28^, indicating that this pair of neurons are DAN-c1 (Figure 2d), thus, this strain will now be referred to simply as DAN-c1.

**Figure 2.**
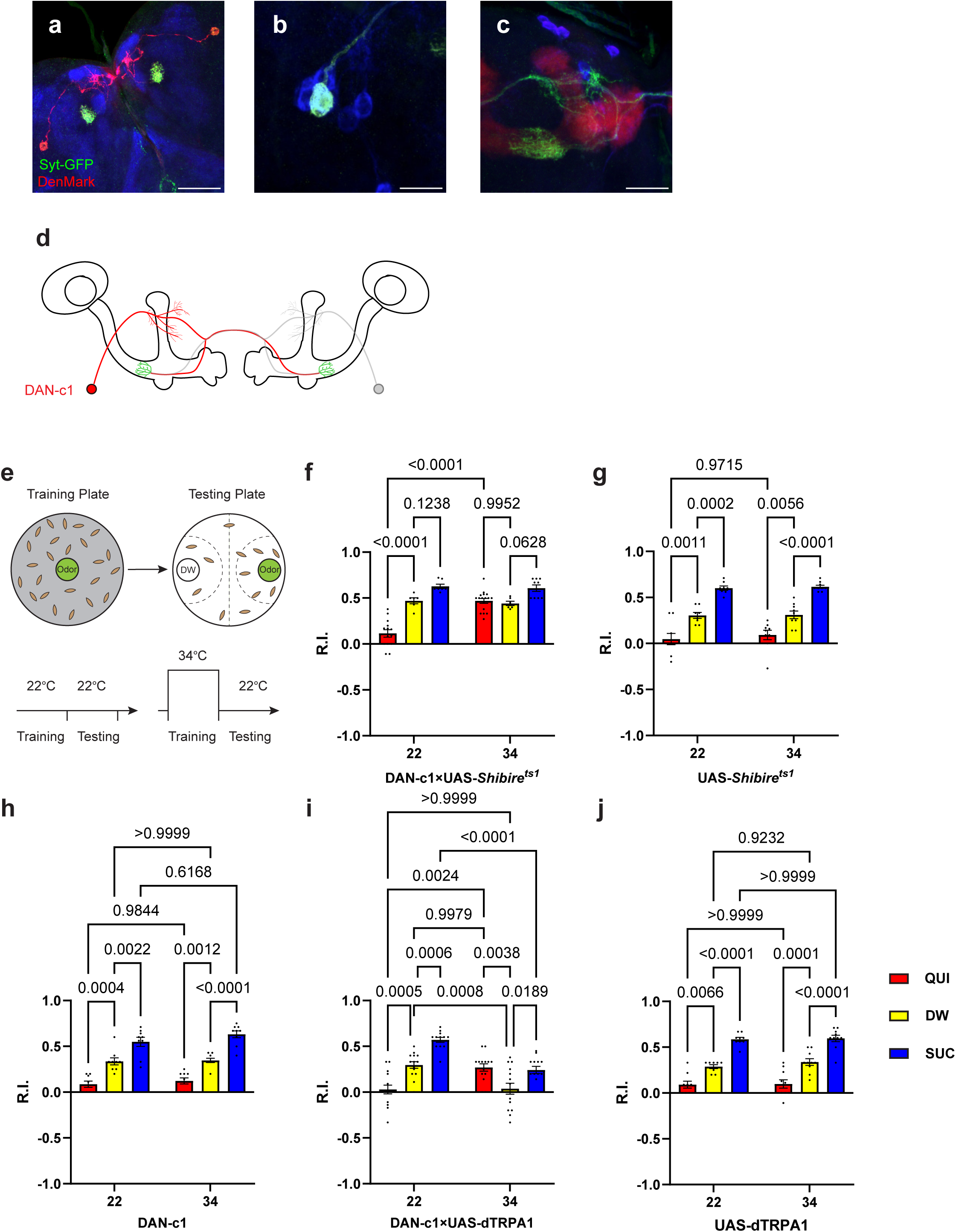
Dopamine release from DAN-c1 mediates larval aversive learning. **(a-d)** R76F02-AD;R55C10-DBD driver is used as it covers one dopaminergic neuron, DAN-c1, in each brain hemisphere. **(a)** Dendrites and axons of DAN-c1 are labeled by DenMark and sytGFP, correspondingly. **(b** and **c)** Soma and neurites from Figure 1f with higher magnification. DAN-c1 is labeled with GFP, the MB with RFP, and DANs with TH antibody (blue color). Only one DAN soma is identified (b), axons from DAN-c1 innervate the lower peduncle (LP) of the MB (c). A schematic diagram (**d**) shows the innervation patterns of DAN-c1. Modified from Eichler *et al.*^44^, and Saumweber *et. al*.^28^. **(e)** A schematic paradigm for larval olfactory learning (top) and two different training paradigms for thermogenetics (bottom). **(f-h)** Blocking dopamine release from DAN-c1 during learning using *shibire^ts1^*strain at 34°C impairs larval aversive learning. **(h-j)** Activation of DAN-c1 with dTRTPA1 at 34°C induces aversive learning. **QUI,** quinine. **DW,** distill water. **SUC,** sucrose. Data are shown as mean ± SEM. Two-way ANOVA, Tukey’s multiple comparison test. For N numbers, interaction p-values, row factor p-values, and column factor p-values, see Table S4. Scale bars: 50 µm (a), 20 µm (b and c). *(Note) N numbers of immunostaining for each strain can be found in Table S2*.

To reveal the role of DAN-c1 in larval olfactory learning, a single odor learning paradigm and thermogenetic tools were applied^17,18,62^. Compared to those trained with distilled water (DW), control strains of larvae exhibited repulsion to the odorant pentyl acetate (PA) after being trained with quinine (QUI) paired with PA, reflecting aversive learning. In contrast, larvae were attracted to PA after being trained with sucrose (SUC) paired with PA, reflecting appetitive learning. The extent of repulsion or attraction was represented with a response index (R.I.) that is compared to the DW group (Figure 2e; For further details, refer to the **Materials and Methods** section).

To examine the role of DAN-c1 in aversive learning, we used a *Shibire^ts1^*strain, which encodes a thermosensitive mutant of dynamin blocking neurotransmitter release when the ambient temperature is higher than 30°C by repressing endocytosis and vesicle recycling functions^17^. When trained with QUI at 34°C, larvae with *Shibire^ts1^* expression in DAN-c1 showed significantly increased R.I. compared to that at room temperature (22°C), while it is not significantly different from the DW at 34°C group (Figure 2f). The complete inactivation of dopamine release from DAN-c1 with *Shibire^ts1^* impaired aversive learning, indicating that dopamine release from DAN-c1 is important for larval aversive learning to occur.

In the next experiments, a fly strain carrying temperature-sensitive cation channel, dTRPA1, was used to excite the DAN-c1 neuron because it can be activated at temperatures higher than 30°C^57^. Compared to those at 22°C, activation of DAN-c1 with dTRPA1 at 34°C during training induced repulsion to PA in the distilled water group, while it is not significantly different from the QUI group at 22°C(Figure 2i). These data suggested that DAN-c1 excitation and presumably increased dopamine release leads to larval aversive learning in the absence of gustatory pairing. Combining the blockade results with *Shibire^ts1^*, these data revealed that dopamine released from DAN-c1 activation mediates larval aversive learning. However, when paired with a gustatory stimulus (QUI or SUC), activation of DAN-c1 during training impairs both aversive and appetitive learning (Figure 2i). We suggest that these data indicate a critical role for the amount of dopamine release from DAN-c1 in larval associative learning, as dTRPA1 stimulation may result in excessive dopamine release (see the **Discussion** section).

### The expression pattern of D2R in the third-instar larval brain

Although dopamine D1-like receptors have been proven important for learning^40^, the role of D2-like receptors has not been fully investigated. In addition, the expression pattern of D2R in fly brains was not reported. A fly strain expressing GFP-tagged D2R (BDSC#60276) was used to reveal the expression pattern of D2R in the third-instar larval brain (Figure 3a). D2Rs were found in dopaminergic neurons and the mushroom body. In dopaminergic neurons (Figure 3b-g), D2Rs were found in DM1, pPAM, DL2b, and some DL1 neurons. In the mushroom body, D2Rs were expressed in the soma and lobes, but not in the calyx (Figure 3h and i). Even though D2Rs were widely found in vertical lobes, medial lobes, and peduncles, they were not expressed in every mushroom body neuron. A transection of the peduncle region showed the absence of D2Rs in the core area (Figure S3i), which is composed of densely packed newly created fibers and lacks Fasciclin II (FAS II)^47^.

**Figure 3.**
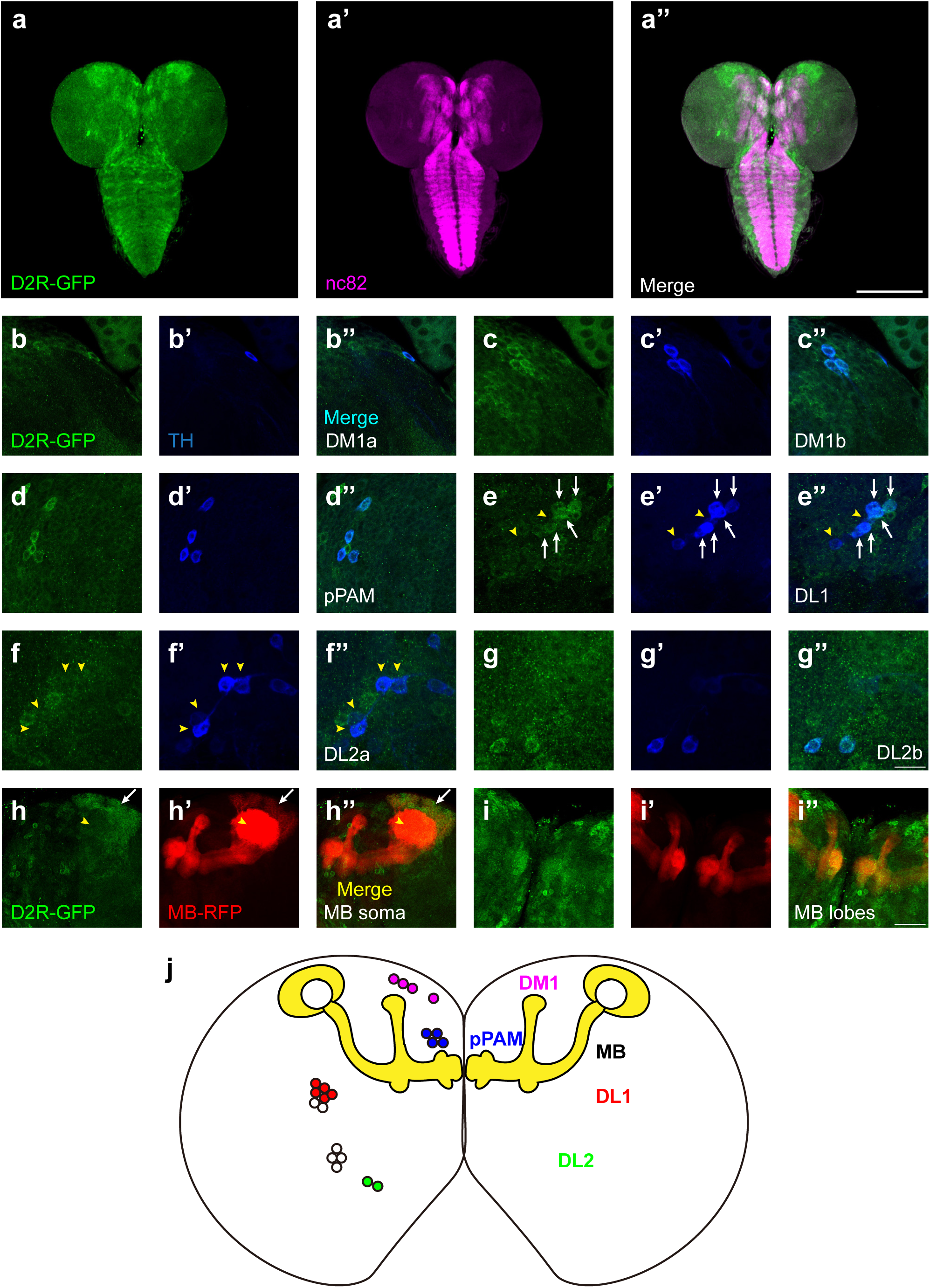
D2Rs are expressed in dopaminergic neurons and mushroom body in *Drosophila* larval brains. **(a)** The expression pattern of D2R in a general view. D2R is shown with tagged GFP (D2R-GFP). Magenta represents neuropils marked by nc82 antibody (a’). **(b-g)** D2Rs (presynaptic) are found in most DANs: DM1a (**b**) and DM1b (**c**), pPAM (**d**), DL1 **(e)**, DL2a **(f)**, and DL2b **(g)** clusters. D2Rs are expressed in parts of DL1 neurons (white arrows in **e**) but not in DL2a neurons (yellow arrow heads in **f**). **(h-i)** D2Rs (postsynaptic) are found in soma of mushroom body (MB) neurons (white arrows in **h**), and MB lobes and peduncles (**i**), but not in calyx (yellow arrow heads in **h**). **(j)** A schematic diagram shows the expression pattern of pre- and postsynaptic D2R in DANs and MB (yellow) in the *Drosophila* larval brain, respectively. Scale bars: 200 µm (a); 50 µm (b-g); 20 µm (h, i). ***Abbreviations:*** DL, dorsolateral; DM, dorsomedial; pPAM, primary protocerebral anterior medial. *(Note) N numbers can be found in Table S2*.

To inspect whether the pattern of GFP signals indeed reflected the expression of D2R, three D2R enhancer driver strains (R72C04, R72C08, and R72D03-GAL4) were crossed with the GFP-tagged D2R strain. R72C08-GAL4 covered three DM1 dopaminergic neurons (Figure S3c), and R72C04-GAL4 labeled one DM1 and two DL2b dopaminergic neurons (Figure S3d and e). R72D03-GAL4 identified parts of mushroom body neurons, whose axons spread on the surface of the mushroom body lobes (Figure S3f); R72C08-GAL4 also identified a subset of neurons from four MB neuroblasts, with soma in four clusters and a converged bundle of axons (Figure S3g and h). These results revealed the expression of D2R in the mushroom body and dopaminergic neurons in the third-instar larval brain.

### D2R in DAN-c1 influences larval aversive learning

Our previous work reported that D2R knockdown (UAS-RNAi) in dopaminergic neurons driven by TH-GAL4 impaired larval aversive learning^62^. Using a microRNA strain (UAS-D2R-miR)^60^, a similar deficit was observed (Figure S4f). To further understand the roles of D2R in aversive learning, its expression in distinct DANs, as well as corresponding learning assays were investigated. Crossing the GFP-tagged D2R strain with a DAN-c1-mCherry strain demonstrated the expression of D2R in DAN-c1 (Figure 4a).

**Figure 4.**
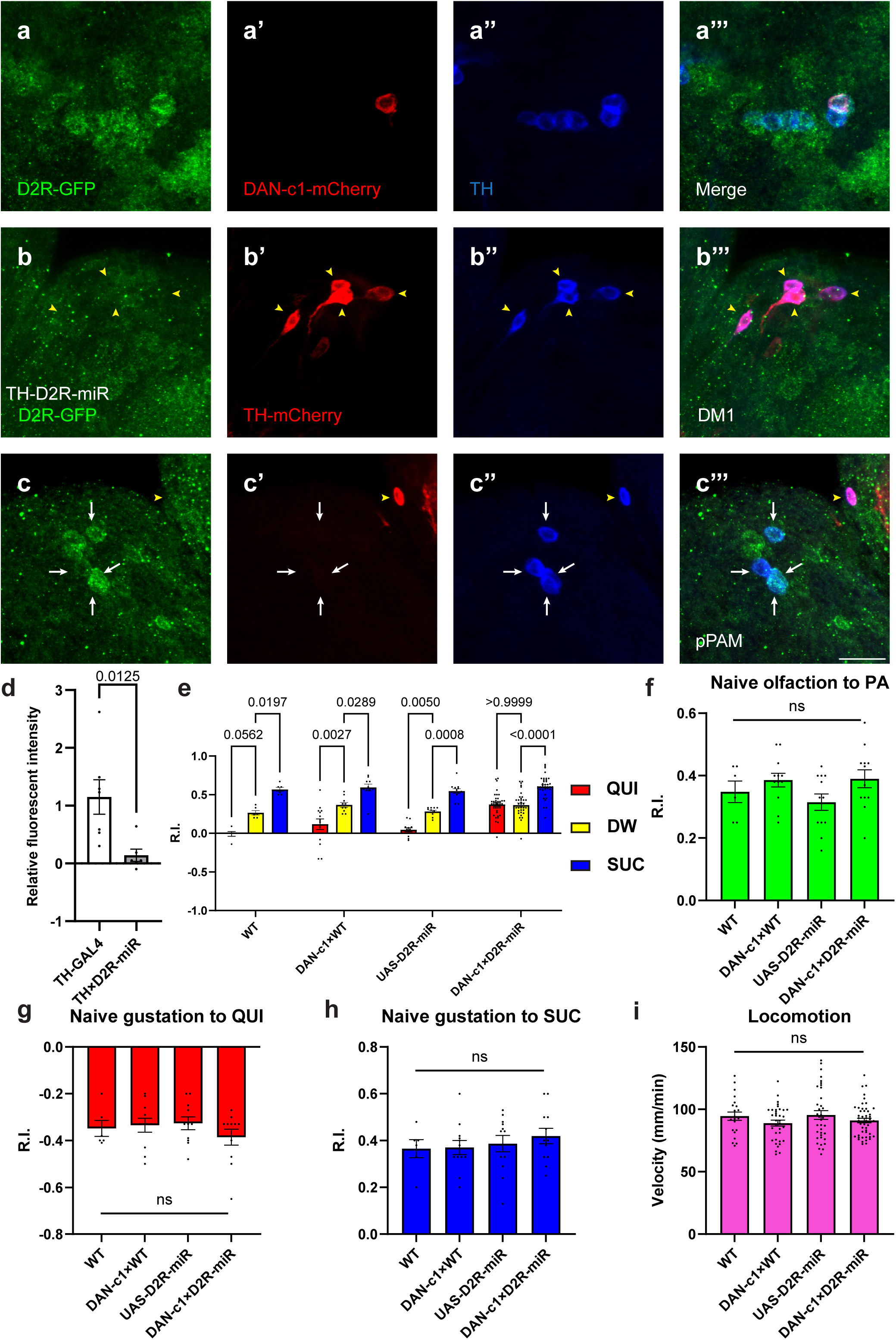
Presynaptic D2R in DAN-c1 is necessary for larval aversive learning. **(a)** D2R is expressed in DAN-c1. The expression pattern of D2R is shown with tagged GFP, DAN-c1 is marked by mCherry, and all DANs are marked with TH antibody (blue). **(b-c)** Knockdown of D2R by D2R-miR reduces fluorescent intensities of D2R-tagged GFP (D2R-GFP). The TH-GAL4 driver is used to express mCherry. The intensity of D2R-tagged GFP is reduced in the DM1 cluster (yellow arrowheads in b), which is still intact in pPAM neurons (white arrows in c). **(d)** D2R-GFP fluorescent intensity is quantified by standardizing the values in DM1 with those in the pPAM. Data are shown as mean ± SEM. For the TH-GAL4 group, N=7 brains; for the TH×D2R-miR group, N=6 brains. Unpaired t-test, p = 0.0125. **(e)** Knockdown of D2R in DAN-c1 by D2R-miR impairs larval aversive learning. **QUI,** quinine. **DW,** distill water. **SUC,** sucrose. Data are shown as mean ± SEM. Two-way ANOVA, Tukey’s multiple comparison test, p < 0.0001 for interaction, row factor (genotype), and column factor (US) p-values. For N numbers, see Table S5. **(f-i)** D2R knockdown in DAN-c1 does not affect naïve sensory and motor functions. Data are shown as mean ± SEM. One-way ANOVA, Tukey’s multiple comparison test. For N numbers, see Table S5. Scale bar: 20µm. *(Note) Figures S5 and S6 show additional information on naïve sensory and motor functions in larvae related to D2R-miR experiments. N numbers of immunostaining for each strain can be found in Table S2*.

To reduce the expression of D2R in dopaminergic neurons, a microRNA strain UAS-D2R-miR was used when crossing with distinct driver strains. The efficiency of D2R knockdown was confirmed by crossing the GFP-tagged D2R strain with TH-GAL4;UAS-D2R-miR. In these larval brains, GFP signals in DM1 were not detected, while those in pPAM were still intact (Figure 4b and c). Quantification showed a significant decrease of GFP signals in the knockdown group compared to the control (Figure 4d), indicating reduced transcripts of D2R linked GFP by D2R-microRNA (Figure S4c).

To investigate the roles of D2R in distinct dopaminergic neurons during larval associative learning, UAS-D2R-miR strain was crossed with distinct driver strains labeling different pairs of dopaminergic neurons. Among them, the knockdown of D2R in DAN-c1 impaired aversive learning with the odorant pentyl acetate, while appetitive learning was unaffected (Figure 4e). In contrast, although D2R was also found in DAN-d1 and DAN-g1, neither D2R knockdown in DAN-d1 nor in DAN-g1 affected larval olfactory learning (Figure S4d-f, see the **Discussion** section). As the naïve sensory and motor functions were not affected, this deficiency was caused by impairment in learning abilities (Figure 4f-i). Similar learning deficits were observed in the same strain trained with another odorant, propionic acid (Figure S5a), as well as in larvae with D2R knockdown using UAS-RNAi (Figure S5b). These results demonstrated that D2Rs are expressed in DAN-c1, and they are necessary for larval aversive learning. Presumably the knockdown of presynaptic inhibitory D2R autoreceptors on DAN-c1 will result in increased and excessive dopamine release, which leads to aversive learning deficiency. These results are consistent with the activation studies with dTRPA1 above, in which increased dopamine release during training results in impaired aversive learning (Figure 2i).

### Over-excitation of DAN-c1 during learning impairs larval aversive learning

To exclude possible chronic effects of D2R knockdown during development, optogenetics was applied at distinct stages of the learning protocol. Channelrhodopsin2 (ChR2) is a blue light activated cation channel from algae, which can be used to activate target neurons^63,64^. Over-excitation of DANs under a TH-GAL4 driver with ChR2 during training impaired aversive learning but left appetitive learning intact (Figure S7), which is consistent with D2R knockdown results. To investigate the mechanisms with a better temporospatial resolution, ChR2 was expressed in DAN-c1, and blue light was applied at distinct stages of the learning protocol (Figure 5a). Optogenetic activation of DAN-c1 during training impaired aversive learning, not appetitive learning (Figure 5b-d). This result is consistent with the effect of D2R knockdown in DAN-c1, indicating that increased, excessive dopamine release during training leads to impaired aversive learning.

**Figure 5.**
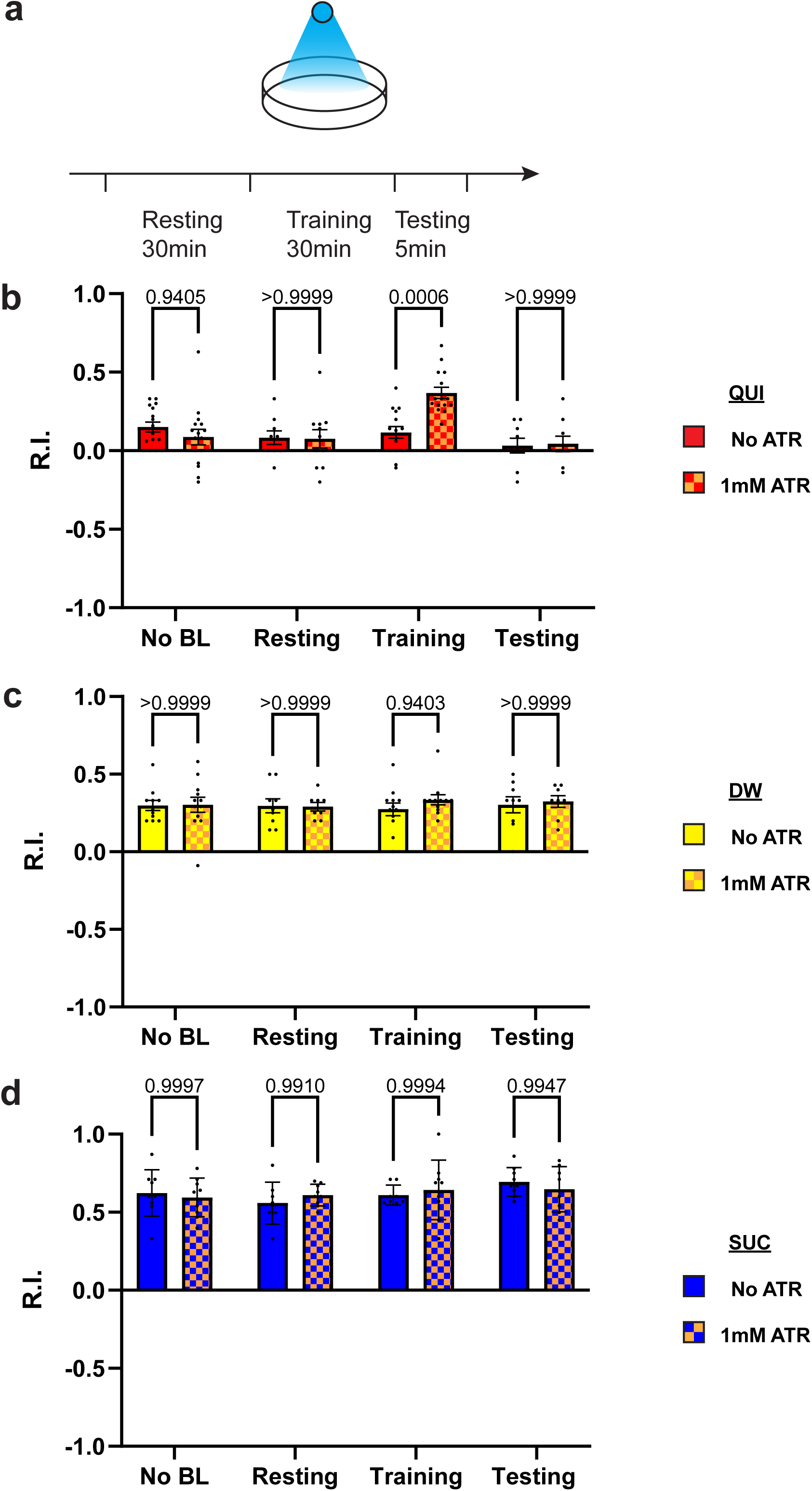
Over-excitation of DAN-c1 impairs larval aversive learning. **(a)** A schematic diagram of optogenetic manipulations of neuronal excitability in DAN-c1 during distinct stages in learning. **(b-d)** Activation of DAN-c1 during training impairs larval aversive learning (**b**), while keeps appetitive learning intact (**d**). Unconditioned stimuli used were quinine (b) and sucrose (d). No learning behaviors are observed in the control distilled water (DW) groups (**c**). Third-instar larvae with ChR2 expression in DAN-c1 are used. *ATR,* all-trans-retinal. Data are shown as mean ± SEM. Two-way ANOVA, Tukey’s multiple comparison test. In QUI group p-values (b), p = 0.0009 for interaction, p < 0.0001 for row factor (training stages), and p = 0.1365 for column factor (whether with ATR); in DW group p-values (c), p = 0.8367 for interaction, p = 0.9750 for row factor (training stages), and p = 0.4872 for column factor (whether with ATR); in SUC group p-values (d), p = 0.6247 for interaction, p = 0.2550 for row factor (training stages), and p = 0.9437 for column factor (whether with ATR). For N numbers, see Table S6.

### D2R in mushroom body mediates larval learning through inhibition

We have shown that D2R in DAN-c1 plays a critical role in larval aversive learning. Since D2Rs are also expressed in soma and axons in most mushroom body neurons (Figure 3h and i), we examined the role of D2R in MB neurons, a center for learning in *Drosophila*. Knockdown of these D2Rs by D2R-miR impaired both appetitive and aversive learning (Figure 6a). Similarly, optogenetic activation of mushroom body neurons during training led to deficiencies in both appetitive and aversive learning (Figure 6b and d). These deficiencies were not observed in larvae with activation during the resting stage. As D2Rs are inhibitory receptors, and optogenetic activation leads to greater neuronal excitation like what may occur with knockdown of D2Rs, these data show that the inhibitory effect of D2Rs in mushroom body neurons is necessary for larval olfactory associative learning to occur.

**Figure 6.**
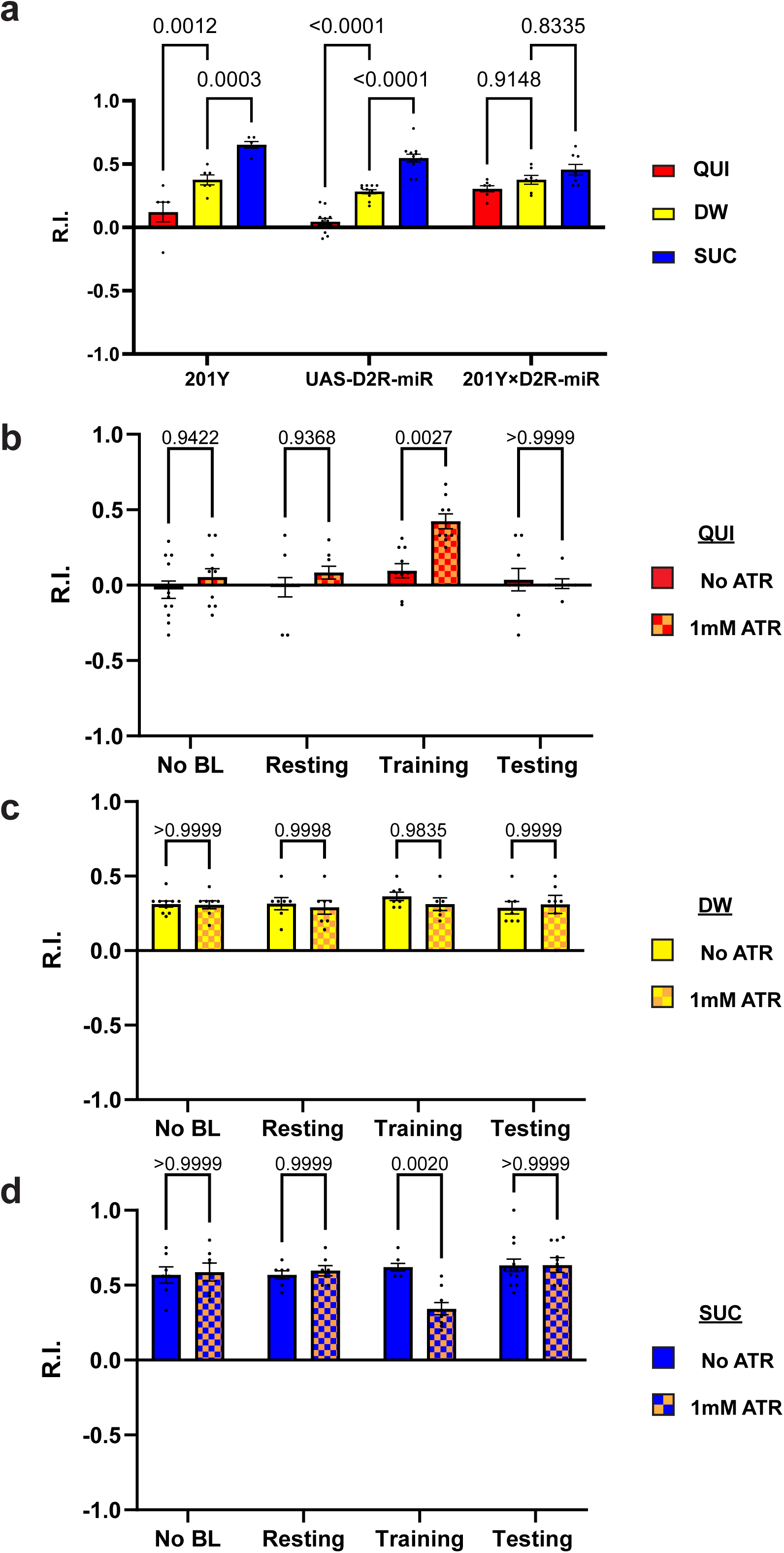
D2R in mushroom body is necessary for both aversive and appetitive learning. **(a)** Knockdown D2R in mushroom body neurons (MBNs) impairs larval aversive and appetitive learning. **(b-d)** Activation of MBNs during training impairs both larval aversive and appetitive learning. Unconditioned stimuli used were quinine (b) and sucrose (d). No learning behaviors are observed in the control distilled water (DW) groups (c). Third-instar larvae with ChR2 expression in MBNs (201Y-GAL4) are used. *ATR,* all-trans-retinal. Data are shown as mean ± SEM. Two-way ANOVA, Tukey’s multiple comparison test. In D2R knockdown experiments (a), p < 0.0001 for interaction, p = 0.0011 for row factor (genotype), and p < 0.0001 for column factor (US). In optogenetic QUI group (b), p = 0.0259 for interaction, p < 0.0001 for row factor (training stages), and p = 0.0040 for column factor (whether with ATR); in DW group (c), p = 0.8077 for interaction, p = 0.7623 for row factor (training stages), and p = 0.5846 for column factor (whether with ATR); in SUC group p-values(d), p = 0.0035 for interaction, p = 0.0086 for row factor (training stages), and p = 0.0865 for column factor (whether with ATR). For N numbers, see Table S5 and S6.

## Discussion

The dopaminergic system plays an important role in *Drosophila* olfactory associative learning, but the roles of D2R in this process have not been fully explored. In this study, we systematically investigated the expression pattern of D2R in the third-instar larval brain as well as its role in larval aversive and appetitive learning. One driver strain identifying a pair of DAN-c1 neurons in the third-instar larval brain was discovered and learning assays with thermogenetic tools (*Shibire^ts1^*, dTRPA1) demonstrated that the blockade of dopamine release from DAN-c1 impeded aversive learning, while its activation during training led to repulsion toward the odor in the absence of unconditioned stimulus (i.e., QUI). These results revealed that DAN-c1 activation (i.e., presumably leading to the release of synaptic dopamine) mediates larval aversive learning to occur. Subsequently, the expression pattern of D2R was explored by using a GFP-tagged D2R strain, including distinct dopaminergic and mushroom body neurons. D2Rs were expressed in DAN-c1, and the knockdown of these receptors resulted in aversive learning deficiency. These data suggested that presynaptic D2Rs in a pair of dopaminergic neurons, DAN-c1, regulate dopamine release during excitation whereas knockdown of these same receptors leads to excessive dopamine release, causing deficits in aversive learning to occur. Furthermore, the activation of DAN-c1 with optogenetic tools during training, resulting in excessive dopamine release, impaired aversive learning, as well. Finally, it was demonstrated that either the knockdown of postsynaptic D2R or activation of mushroom body neurons led to learning deficits. These data demonstrate that D2Rs in distinct brain locations are critically involved in associative learning.

### The characteristics of the single odor larval learning paradigm

We adopted the single odor larval learning paradigm from previous publications^17,18^. To validate this paradigm induces associative learning responses, Honjo et al.^17,18^ tested the paradigm from four aspects: First, the paradigm did not show obvious sensitization or habituation effects when larvae were tested. They applied the odorant to the larvae after training, only the ones had paired training with both odor and unconditioned stimulus (quinine or sucrose) showed learning responses. Larvae exposed for 30 minutes to either the odorant or the unconditioned stimulus alone did not show a different response to the odor compared to the naïve group. Second, the odor responses are associative. Honjo et al. showed only when the odorant was paired with unconditioned stimulus would induce corresponding attraction or repulsion of larvae to the odor. Neither odorant alone, unconditioned stimulus alone, nor temporal dissociation of odorant and unconditioned stimulus would induce learning responses. Third, the odor responses are odor specific. When applied to a second odorant that was not used for training, larvae only showed learning responses to the odor paired with unconditioned stimulus. This result ruled out the explanation of a general olfactory suppression and indicated larvae can discriminate and specifically alter the responses to the odor paired with unconditioned stimulus. Although the two-odor reciprocal training is not used, these results can show the association of unconditioned stimulus and the corresponding paired odor. Finally, well known learning deficit mutants did not show learned responses in this learning paradigm. Honjo et al. tested mutants (e.g., *rut* and *dnc*), which showed learning deficits in the adult stage with two odor reciprocal learning paradigm^21,25^. These mutant larvae also failed to show learning responses when tested with the single odor larval learning paradigm. Combining all the evidence above, we believe this single odor larval learning paradigm is a robust and reliable paradigm for larval associative learning assays, composing essential characteristics of classical conditioning. Previously, we applied this paradigm to investigate the roles of mushroom body serotonin receptors (5-HT7) in larval appetitive learning^65^. In this study, we used two distinct odorants (pentyl acetate and propionic acid), as well as two D2R knockdown strains (UAS-miR and UAS-RNAi for D2R). We obtained similar results for larvae with D2R knockdown in DAN-c1 using different odorants or D2R knockdown strains. In addition, our naïve olfactory, naïve gustatory, and locomotion data ruled out the possibilities that the responses were caused by impaired sensory or motor functions. Comparison with the control group (odor paired with distilled water) ruled out the potential effects if habituation existed. All these results support this single odor learning paradigm is reliable to assess the learning abilities of *Drosophila* larvae. The failure of reduction in R.I. when larvae with D2R knockdown in DAN-c1 were trained in quinine paired with the odorant is caused by deficit in aversive learning ability.

### Insights into the neuronal circuits underlying larval olfactory associative learning

Mushroom body and dopaminergic neurons play important roles in *Drosophila* associative learning in both larvae^66^ and adults^66–69^. Combining our results with previous learning circuitry research in adult flies and larvae, we hypothesized the mechanism underlying larval olfactory associative learning. Olfactory information (odors, <u>C</u>onditioned <u>S</u>timulus, CS) is received by olfactory sensory neurons (OSN), then transmitted to the mushroom body via projection neurons (PN)^70^. The mushroom body (MB) is a primary learning center of *Drosophila* and composed of Kenyon cells (KC, or mushroom body neurons, MBN)^71–73^. Their dendrites form the calyx, receiving olfactory information from projection neurons. The axons converge into peduncles, then branch into the vertical and medial lobes. Distinct gustatory cues (taste, <u>U</u>nconditioned <u>S</u>timuli, US) are sensed by gustatory sensory neurons (GSN) and transferred to dopaminergic neurons in different clusters (Figure 7a). DAN-c1 in the DL1 cluster mediates aversive cues, projects to the lower peduncle (LP) in the mushroom body. The plasticity of synapses from MBNs to MB output neurons (MBON) may be negatively modulated by dopamine, like those in adults^67,74,75^. The MBN-MBON synapses in the vertical lobe and peduncle are responsible for attraction, while those in the medial lobe are for repulsion.

**Figure 7.**
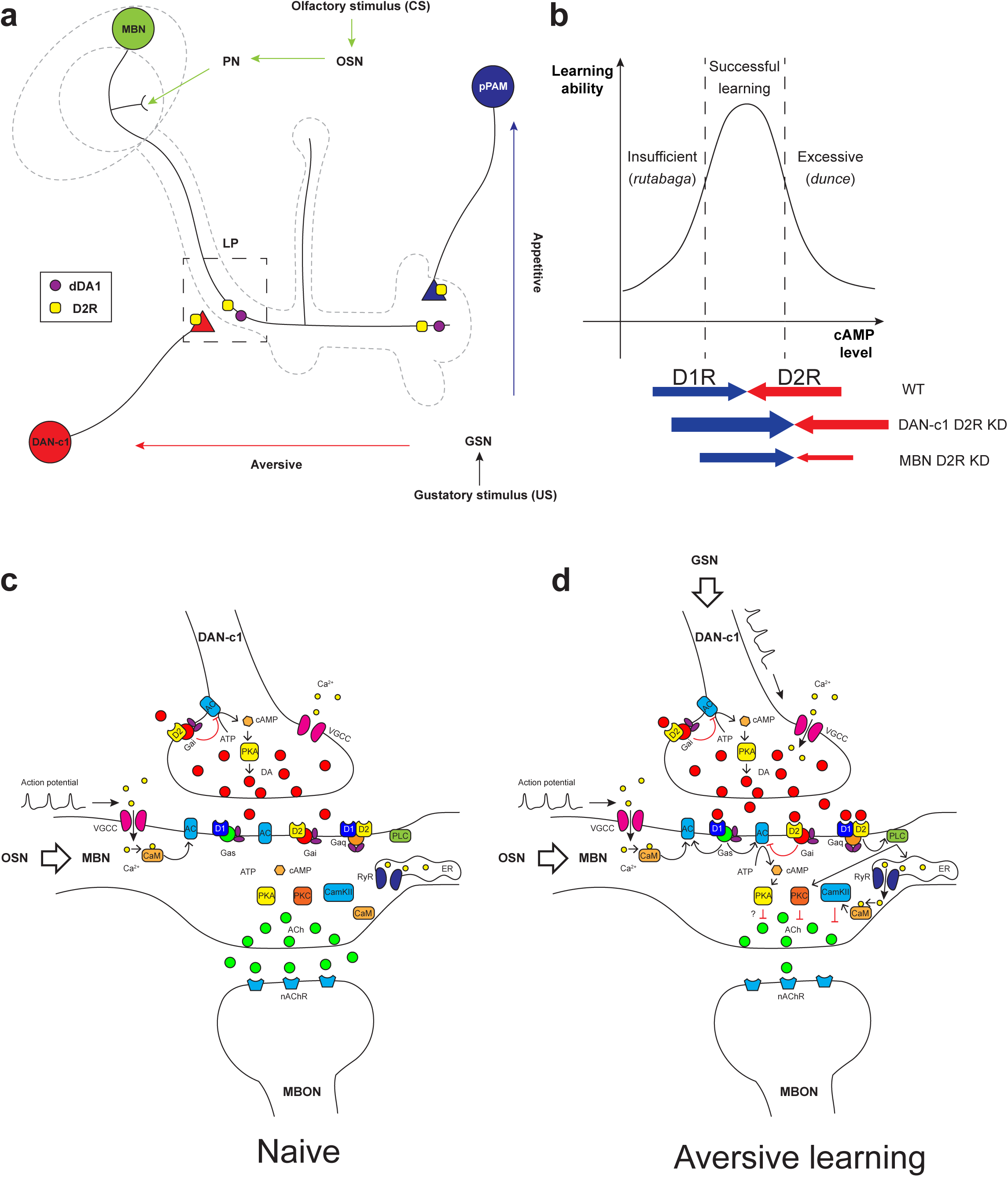
Roles of D2Rs in dopaminergic and mushroom body neurons during larval olfactory learning. **(a)** A schematic diagram shows the roles of D2R in dopaminergic neurons (DAN) and mushroom body neurons (MBN) in larval olfactory associative learning. During learning, olfactory stimuli (conditioned stimulus, CS) are received by olfactory sensory neurons (OSN) and transmitted to MBNs (green) via projection neurons (PN). Distinct gustatory stimuli (unconditioned stimulus, US) are received by gustatory sensory neurons (GSN) and transferred to different dopaminergic neurons. Aversive stimuli are sent to DAN-c1 (red) in the DL1 cluster which connects the lower peduncle compartment (LP, square in dash line), while appetitive stimuli are received by pPAM neurons (blue) innervating the medial lobe (ML). D2Rs (yellow square) are expressed in DAN-c1 and pPAM as autoreceptors, regulating dopamine release. Both D2R and dDA1 (magenta circle) are expressed in the MBNs. **(b)** A hypothetical curve showing the relationship between learning ability and cAMP level in the mushroom body. Insufficient cAMP (*rutabaga mutant*) cannot induce learning, while excessive cAMP (*dunce mutant*) also impairs learning. Only the appropriate level of cAMP regulated by the opposing actions of D1R (dDA1) and D2R leads to successful learning in wild type larvae (WT). Knockdown of D2R in DAN-c1 causes excessive dopamine release, elevating cAMP and resulting in impaired learning. D2R knockdown in MBNs relieves the inhibition effect of D2R, resulting in excessive intracellular cAMP and learning failure. **(c-d)** Potential molecular mechanisms underlying *Drosophila* olfactory learning in the square region shown in (a). **(d)** During aversive learning, olfactory stimuli induce depolarization of MBNs, which activates voltage-gated calcium channels and induces calcium influx. Gustatory stimuli, such as quinine, activates dopaminergic neurons and elevates dopamine (DA) release. D1 receptor (dDA1) activates adenylyl cyclase (AC) and elevates cAMP via Gα_s_, while D2 receptor (D2R) inhibits AC and suppresses cAMP via Gα_i/o_. In *Drosophila*, the coincidence detector *rutabaga* (AC) is activated by the existence of both calcium and Gα_s_, converging the olfactory and gustatory stimuli. cAMP activates the PKA signaling pathway, elevating the neuronal excitability. D1 and D2 receptors can also form heteromeric receptors and activate the PLC-PKC and CaMKII signaling pathways via Gα_q_. These pathways inhibit acetylcholine (ACh) release from MBNs to MB output neurons (MBON), which leads to avoidance of the learned odor. ***Abbreviations:*** AC, adenylyl cyclase; ACh, acetylcholine; ATP, adenosine triphosphate; CaM, calmodulin; CaMKII, Ca^2+^/calmodulin-dependent protein kinase II; cAMP, cyclic adenosine monophosphate; CS, conditioned stimulus; DA, dopamine; DAN, dopaminergic neurons; DL, dorsolateral; ER, endoplasmic reticulum; Gα_i/o_, G_i/o_ protein α subunit; Gα_q_, G_q_ protein α subunit; Gα_s_, G_s_ protein α subunit; GSN, gustatory sensory neurons; LP, lower peduncle; MBN, mushroom body neurons; MBON, mushroom body output neurons; ML, medial lobe; nAChR, nicotinic acetylcholine receptor; OSN, olfactory sensory neurons; PKA, protein kinase A; PKC, protein kinase C; PLC, phospholipase C; PN, projection neurons; pPAM, primary protocerebral anterior medial; RyR, ryanodine receptor; US, unconditioned stimulus; VGCC, voltage gated calcium channel.

When only the odorant appears, the subset of Kenyon cells representing this odor may be depolarized, inducing calcium influx as in adult flies^73^. As a balance exists between compartments across the mushroom body lobes, the response to the odor depends on the naïve olfactory circuits from projection neurons to the lateral horn (LH). In aversive learning, in addition to olfaction induced calcium influx, gustatory stimuli also lead to dopamine release from DAN-c1 and subsequently, activation of Gα_s_ in LP. The co-existence of calcium and Gα_s_ activates a Ca^2+^-dependent adenylate cyclase (AC), rutabaga (rut, as in adults^76–79^). Rutabaga converges information from both olfaction and gustation, working as the coincidence detector of associative learning. Its downstream signaling inhibits attractive MBN-MBON synapses in the LP, breaking the balance between distinct compartments. As a result, after learning, the larvae will exhibit repulsion when exposed to the odor again. In contrast, dopaminergic neurons in the pPAM convey appetitive cues, which project to compartments in the medial lobe. The co-existence of olfactory and appetitive gustatory stimuli leads to inhibition of these repulsive MBN-MBON synapses, inducing attraction.

### The conserved role of DAN-c1 in aversive learning throughout *Drosophila* development

Adult *Drosophila* share similar neuronal circuits of learning with larvae^28,53,66–69^. In adult brains, dopaminergic neurons are classified into 13 clusters, named as PAM (protocerebral anterior medial), PAL (protocerebral anterior lateral), PPM (protocerebral posterior medial), PPL (protocerebral lateral), and PPD (protocerebral posterior dorsal) clusters^80^. DANs in the PAM cluster innervate the medial lobe, while those in PPL1 project to the vertical lobe^69^.

R76F02-AD;R55C10-DBD identifies two dopaminergic neurons in adult brains, MB-MP1 in PPL1 and ALT-PLPC in PPL2ab^60^. MB-MP1 is also named PPL1-γ1pedc, innervating both γ1 and the peduncle of the β lobe^69^. Activation of this neuron induced aversive learning^61^, and activation of its corresponding MBON-γ1pedc>α/β led to approach^81^. During metamorphosis, dopaminergic neurons in DL1 develop into the PPL1 cluster, DL2a neurons develop into PPL2ab, and those in pPAM will develop into the PAM cluster^82^. This driver strain only identifies DAN-c1 from DL1 in larvae, which innervates the lower peduncle of the mushroom body. Interestingly, previous reports revealed that memory can be transferred from larvae to adults^83^, indicating the maintenance of neuronal circuitry architecture during metamorphosis. This evidence supports that DAN-c1 is the corresponding neuron of PPL1-γ1pedc in larvae, which performs similar functions in aversive learning.

### Pre- and post-synaptic D2Rs regulate cAMP in the mushroom body during aversive learning

The molecular mechanisms underlying *Drosophila* learning have not been fully determined. In a traditional view, gustatory cues elevate dopamine release, which binds to D1 receptors and then activates Gα_s_. The co-existence of Gα_s_ and calcium elicited by olfactory cues activates rutabaga in axons of MBNs. Rutabaga transforms ATP into cAMP, activating PKA signaling pathway. Mutant flies with either insufficient (*rutabaga*) or excessive cAMP (*dunce*) showed aversive learning deficiency, indicating that the level of cAMP should be kept in an optimal range to achieve aversive learning^21,25^ (Figure 7b).

Our results have shown that D2Rs in dopaminergic neurons and the mushroom body are important for larval aversive learning, suggesting a “*dual brake*” role in regulating cAMP levels in the MBNs through both pre- and postsynaptic components. On the presynaptic side, D2R in DAN-c1 decreases the release of dopamine under gustatory stimuli, reducing the probability of postsynaptic D1R activation in MBNs. On the postsynaptic side, D2R in MBNs inhibits the coincidence detector rutabaga (AC) both via activation of Gα_i/o_ and inhibition of voltage-gated calcium channels^2^, indicating postsynaptic D2R functioning as a “*brake of the coincidence detector”*. Combining these, “*dual brake*” D2R ultimately regulates the mushroom body cAMP level within a physiologically optimal range during aversive learning. In addition, D2R fine tunes the functional concentration spectrum of dopamine with higher resolutions, as its dopamine affinity is 10- to 100-fold greater than D1 receptors. Overall, D2Rs work in a “*dual brake*” system both expanding the representation of a dynamic range of gustatory signal intensity with high signal to noise ratio and preventing postsynaptic overexcitation, which increases the reliability of DAN-c1 and MBN circuits for the larval aversive learning (Figure 7c and d).

Recent studies showed that the approach/repulsion in adult learning is achieved via inhibition of the repulsive/attractive representing compartments^67,74,75^, which indicates dopamine inhibits acetylcholine release from MBN to MBON^84^. However, the PKA signaling pathway usually elevates neuronal excitability and increases neurotransmitter release^2,3^, which is contradictory to the recent findings^85^. In addition to the “*dual brake*” role of D2R, our results suggest a third role of D2R in aversive learning. D1 and D2 receptors can form heteromeric receptors, whose downstream Gα_q_ activates PKC and CaMKII signaling pathways. The activation of these signaling pathways may reduce acetylcholine release^86^ (Figure 7c and d).

### Explanation of the results of thermogenetic and optogenetic experiments

Activation of DAN-c1 with dTRPA1 induced aversive learning, while the repulsion disappeared when DAN-c1 was activated in the quinine group (Figure 2i). Our explanation is that quinine stimulation and temperature activation led to over-excitation of DAN-c1, which impaired aversive learning. This is consistent with the learning deficiency in larvae with D2R knockdown in DAN-c1 (Figure 4e). Results from optogenetic activation of DAN-c1 during aversive learning also support this (Figure 5b). However, in contrast to results with thermo-activation, larvae with optogenetic activation of DAN-c1 showed neither repulsion after being trained with distilled water (Figure 5c), nor did they show reduced attraction in the sucrose group (Figure 5d). One possible explanation is that the thermo-activation is relatively mild compared to the optogenetic activation. Based on this, thermo-activation of DAN-c1 is still in the physiological range of cAMP under the upper limit (Figure 7b), resulting in repulsion in the distilled water (DW) group, neutralized attraction in SUC group, and impaired repulsion in QUI group. In contrast, optogenetic activation of DAN-c1 overwhelmed the physiological conditions, leading to failure of repulsion in DW group (Figure 5c). This repulsive failure did not affect appetitive learning (Figure 5d), and a stronger over-excitation in QUI group induced similar failure (Figure 5b).

### Distinct dopaminergic neurons may have different roles in larval aversive learning

Although D2Rs are also expressed in DAN-d1 and DAN-g1 (Figure S4d and e), the knockdown of D2R in these neurons did not impair larval aversive learning (Figure S4f). For DAN-g1, interestingly, the R.I. from D2R knockdown larvae trained with quinine (QUI, aversive learning) showed significant difference when compared to the DW (control) group, but it was also significant different from the DAN-g1 genetic control group trained with QUI (two-way ANOVA, Tukey’s multiple comparisons, p=0.0002), while not significant different from UAS-D2R-miR genetic control group trained with QUI (p=0.2724). Besides, D2R knockdown in DAN-g1 when trained with another odorant propionic acid (ProA) did not show aversive learning deficiency (Figure S5a). In addition, knockdown D2R in DAN-g1 using RNAi also did not show aversive learning deficiency when trained with odorant pentyl acetate (PA, Figure S5b). This discrepancy may be caused by the different stimulus intensity of distinct odorants, as well as the different efficiency of distinct knockdown methods (microRNA or RNAi strains we used). We supposed that D2Rs in DAN-g1 may partially affect larval aversive learning in a quantitative level; but not play an important role as those in DAN-c1, which will cause a qualitative change when knockdown.

Previous work reported that aversive olfactory learning was induced through the optogenetic activation of DAN-d1, f1, or g1, but not DAN-c1^87^. This discrepancy can be explained as the optogenetic overexcitation of DAN-c1, similar to our optogenetic or D2R knockdown results. Our learning assay results from larvae with D2R knockdown in DAN-d1 or g1 also supported this: aversive learning was not affected by D2R knockdown (Figure S4f). These data indicate D2Rs in DAN-d1 or g1 may not be important in larval aversive learning. Additionally, the DAN-c1 strain used in the previous work (SS02160-split-GAL4) not only labels DAN-c1, but also marks other non-dopaminergic neurons, which may affect the results. Besides, live calcium imaging showed that DAN-d1, f1, and g1 responded to the activation of mechanosensory and nociceptive neurons^87^, indicating a functional differentiation from gustatory activated DAN-c1^61,88^.

In future studies, the molecular signaling downstream of D2R needs to be explored, as well as the comprehensive neuronal circuit architectures of larval learning. The neuronal circuits underlying learning and memory are complex networks, sharing similarities with the regulatory networks of gene expression. Studies of the mechanisms of learning and memory help us understand the essential principles of the non-linear dynamic characteristics in these complex systems. On one hand, the progress in larval learning provides useful information for helping our understanding about more complex systems, from brains in adult flies to those in mammals. On the other hand, the architecture of larval learning circuits could improve either hardware design or algorithm structures in artificial intelligence, which may bring more powerful tools, such as navigation systems regulating multiple auto-drive vehicles in complex 3-dimentional environments.

In conclusion, we explored the expression pattern of D2R in the third-instar larval brain and investigated their roles during larval olfactory learning. D2Rs were found in DAN-c1, and their knockdown induced deficiency in aversive learning. D2Rs were also expressed in MBNs, knockdown of which impaired both aversive and appetitive learning. This research revealed the important roles of D2Rs in *Drosophila* larval olfactory learning and enriched our understanding regarding the mechanisms underlying the learning process.

## Materials and Methods

### Fly stocks

All fly strains used in this study are listed in Table 1 and Table S1. Flies were maintained on standard medium, which consists of cornmeal, yeast, dextrose, sucrose and agar in water. Flies were kept in a 12/12-hour light/dark cycle at 25℃. Canton-S genotype (WT) and yw^1118^ were used as wild-type. Strains carrying more than one transgene were constructed by standard genetic crosses with the w^1118^; CyO/Sco; TM2/TM6 multiple balancer chromosome strain. Strain UAS-Syb::spGFP1-10, LexAop-CD4::GFP11, LexAop-rCD2::RFP /CyO; MB247-LexA::Up16/TM6B was made by chromosome swapping between UAS-Syb::spGFP1-10, LexAop-CD4::GFP11/CyO (II, second chromosome) and LexAop-rCD2::RFP (II).

The D2R gene is located on the X chromosome, with 6 different alternative splicing products. The GFP-tagged D2R strain is inserted with a GFP gene in the second intron, generating D2R molecules tagged with GFP^89,90^. The D2R-miR strain produces microRNA recognizing the sequence across the third and fourth exons, which is not affected by the GFP insertion (Figure S4c).

### GRASP

Split-GFP reconstitution across synaptic partners (GRASP) was used to investigate whether neurons formed synapses^50^ (Figure S2d). Half of split GFP tethered to the presynaptic synaptobrevin (UAS-syb::spGFP1-10) was expressed in one type of neuron using UAS/Gal4 binary system, and the complementary split GFP linked to a membrane protein (LexAop-CD4::spGFP11) was expressed in another category of neuron with LexA/LexAop. If synapses between these neurons exist, the split GFPs would form a complete one and be recognized by a mouse antibody (Figure S2a-c).

### Immunohistochemistry

All staining processes were performed in 1.5 ml Eppendorf tubes. Late third-instar (96-100h after egg laying) larval brains were dissected in dissection solution (300 mOsmol/L). Brains were fixed in 4% paraformaldehyde (PFA, Electron Microcopy Sciences, Cat. No. 15713) for 1 hour on ice. After three washes (0.1% bovine serum albumin in 10mM PBS, Sigma Life Sciences A9647), brains were incubated in the blocking and permeabilization solution (0.2% Triton X-100 in 10mM PBS with 5% normal goat serum; Triton-X 100, Sigma, T8532; NGS, Sigma-Aldrich, G9023) for 2 hours at room temperature. Incubation with primary antibodies was done overnight at 4°C. After three washes, brains were incubated in the secondary antibodies for two hours. Both the primary and secondary antibodies were diluted in the blocking and permeabilization solution. GFP antibody (Rabbit, Thermo Fisher Scientific, Cat. No. A6455, 1:1000), TH antibody (Mouse, Immunostar, Cat. No. 22941, 1:1000), goat anti-rabbit with green fluorescence (Invitrogen, Alexa Fluor 488 conjugate, Cat. No. A-11035, 1:1000), and goat anti-mouse IgG with far-red fluorescence (Alexa Fluor 633 conjugate, Cat. No. 21052, 1:500) secondary antibodies were used. After three washes, brains were transferred and mounted in the Fluoro-Gel with Tris Buffer (Electron Microcopy Sciences, Cat. No. 17985-10) on a piece of micro cover glasses (Electron Microcopy Sciences, Cat. No. 72200-41). Finally, the samples were covered with another piece of micro cover glasses.

A seven-day staining protocol was used for staining GFP-tagged D2R or GRASP, in which brains were fixed in 1% paraformaldehyde in Schneider’s insect medium (Sigma Life Sciences, Cat.NO. S0146) overnight at 4°C. In the second day, the brains were rinsed and washed twice with PAT3 solution (0.5% Triton X-100 in 10mM PBS with 0.5% BSA), each for 1 hour. Then brains were incubated in the blocking and permeabilization solution (3% NGS in PAT3) for 2 hours at room temperature. Incubation of primary antibodies was done overnight at 4°C. On the third day, brains were rinsed and washed twice with PBT solution, then incubated in the secondary antibodies for five days. Both the primary and secondary antibodies were diluted in the PBTN solution. In the staining of GFP-tagged D2R, GFP antibody (Rabbit, ThermoFisher Scientific, Cat. No. A6455, 1: 1000), TH antibody (Mouse, Immunostar, Cat. No. 22941, 1:1000), and mCherry antibody (Rat, ThermoFisher Scientific, Cat. No. M11217, 1:1000) were used. For staining of GRASP, GFP antibody (Mouse, ThermoFisher Scientific, Cat. No. G6539, 1:100) was used. Goat anti-rabbit with green fluorescence (Invitrogen, Alexa Fluor 488 conjugate, Cat. No. A-11035, 1:1000), goat anti-mouse with green fluorescence (Alexa Fluor 488 conjugate, Cat. No. A11029, 1:1000), goat anti-mouse with far-red fluorescence (Alexa Fluor 633 conjugate, Cat. No. A21052, 1:500), and goat anti-rat with red fluorescence (Alexa Fluor 546 conjugate, Cat. No. A-11081, 1:1000) secondary antibodies were used. In the seventh day, brains were rinsed and washed twice with PBT solution, and then transferred and mounted in the Fluoro-Gel with Tris Buffer on a piece of micro cover glasses. Finally, the samples were covered with another piece of micro cover glasses.

### Confocal imaging

All images were obtained with a Zeiss Laser Scanning Microscope 510 (LSM510, Carl Zeiss, Inc., USA). Under 40× and 100× objective magnifications, images were collapsed from confocal stacks of 1.0 μm optical slices. Under 25× objective magnifications, images were collapsed from confocal stacks of 2.0 μm optical slices. Under 10× objective magnifications, images were collapsed from confocal stacks of 12.5 μm optical slices. ImageJ software was used to remove other signals outside of the mushroom bodies, as the background noise in GRASP is strong.

### Larval olfactory learning assays

All control strains used in learning assays were homozygous except DAN-c1×WT, while all experimental groups (D2R knockdown, thermogenetics, and optogenetics) used were heterozygous by crossing the corresponding control strains. The single odor learning paradigm was slightly modified from previous publication^17,18^. In brief, 25 to 50 third-instar larvae (92–96 h after egg laying) were trained on a 2.5% agar plate (100 mm petri dish) covered with 2 mL of 1 M sucrose solution (SUC, Sigma, Cat. No. S1888) or 0.1% quinine hemisulfate solution (QUI, Sigma, Cat. No. 22640). Distilled water (DW) was used as a control. An odorant pentyl acetate (PA, 10μL, Sigma-Aldrich, CAT. No. 109584) was placed on a small piece of filter paper (0.25cm^2^ square) inside the lid. After 30 minutes, larvae were rinsed and transferred to the middle line of a new 2.5% agar plate. A small piece of filter paper (0.25cm^2^ square) with 2.5μL pentyl acetate was placed on one side of the plate, while distilled water on the other side. After 5 min, the numbers of larvae in the two semicircular areas were counted and the response index (R.I.) was calculated with the following equation (Figure 2e):

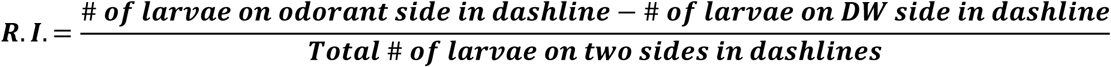

### Naïve Olfactory Test

Larvae were transferred into the midline of test plates. 2.5μL of odorant were added on a piece of filter paper (0.25cm^2^ square) on one side and distilled water on the other side. The number of larvae in two semicircular areas were counted and the R.I. was calculated after 5 min.

### Naïve Gustatory Test

A petri dish with a median separator was used. Both sides were filled with 1% agar, with 2 mL of distilled water on the control side, and with 1 M sucrose (SUC) solution, or 0.1% quinine hemisulfate (QUI) solution on the test side. Twenty larvae were put on each side near the midline and allowed to move for 5 min. Gustatory R.I. was calculated using the larvae numbers on two sides^91^.

### Larval locomotion assay

Individual larvae were placed on the surface of a plate of 2.5% agar mixed with 1mL India ink. They were allowed to acclimate for 1 min, and then a video was recorded for 30 seconds using a Moticam3 digital camera (Motic) and Motic Images Plus 2.0 software. The video was analyzed by the MTrack2 plug-in (from http://valelab.ucsf.edu/∼nico/IJplugins/MTrack2.html) in ImageJ. The path was recorded; scores were quantified as the length traveled per minute as previously described^92^. As the locomotion speed of DAN-c1 homozygous was slow, DAN-c1 × WT was used as the control group.

### Learning assays with thermogenetics

In learning assays with thermogenetics, 25 to 50 third-instar larvae (92–96 h after egg laying) were trained on a 2.5% agar plate (100mm petri dish) covered with 2 mL of 1 M SUC or 0.1% QUI. DW was used as a control. An odorant PA was placed on a small piece of filter paper (0.25cm^2^ square) inside the lid. Training plates were put in a water bath either under 22°C or 34°C. After 30 minutes, larvae were rinsed and transferred to the testing plate. After 5 min, the response index (R.I.) was calculated.

### Learning assays with optogenetics

In learning assays with optogenetics, egg laying plates with 1mM ATR (all-trans retinal, Sigma, Cat. No. R2500) were used. All-trans retinal (ATR) is a necessary light-isomerizable chromophore for ChR2, which is not synthesized by *Drosophila*.^93,94^ Around 50 third-instar larvae were trained in a 35mm petri dish with 2 mL of either 1 M SUC or 0.1% QH solutions. DW was used as a control. During training, an odorant was placed on a small piece of filter paper (0.25cm^2^ square) inside the lid. To activate channelrhodopsin2, a LED (Luxeon Rebel Color LEDs, 07040 PB000-D, wavelength 470 nm) with a power supply (GW Instek, Laboratory DC power supply Model GPS-1830D) was used. The intensity of the blue light was 25 mW, measured by a laser power meter (Sanwa, LP1). After being trained for 30 minutes, larvae were rinsed and transferred to the middle line of a 2.5% agar plate in 100 mm test plate. A small piece of filter paper (0.25cm^2^ square) with pentyl acetate was placed on one side of the plate, while distilled water on the other side. Then the number of larvae in the two semicircular areas were counted and the R.I. was calculated after 5 min.

### Quantification of D2R knockdown

Quantification of the fluorescent intensity of D2R knockdown was performed as follows. TH signals were used to define dopaminergic neurons, and the mean fluorescent intensity of GFP in each neuron was calculated with subtraction of the background. The mean intensity of DM1 was divided by that of pPAM in each brain, and the value in the knockdown group was subsequently normalized with the control group.

### Statistical Analysis

Information for statistical analysis is provided in figure legends. In brief, two-way ANOVA and Tukey’s multiple comparison test were used in Figure 2f-j, Figure 4e, Figure 5b-d, Figure 6a-d, and Figure S7; unpaired t-test was used in Figure 4d; one-way ANOVA and Tukey multiple’s comparison test were used in Figure 4f-i; two-way ANOVA and Dunnett’s multiple comparison test were used in Figure S4f, and Figure S5a-b; one-way ANOVA and Dunnett’s multiple comparison test were used in Figure S6a-i.

## Author contributions

**Qi. C.:** Conceptualization, Methodology, Formal analysis, Investigation, Writing – Original Draft, Visualization, **Qian. C.:** Writing – Review & Editing, **Steijvers. E.:** Writing – Review & Editing **Colvin. R.:** Writing – Review & Editing **Lee. D.:** Conceptualization, Writing – Review & Editing, Supervision, Funding acquisition

## Acknowledgements

This work was partially supported by an NIH grant (AG065925) and an International Collaboration Grant from Korea Institute of Science & Technology (Brain Science Institute), Seoul, Korea. CQ was a recipient of the SEA and GSR awards from Ohio University. We thank Dr. J Hirsh (University of Virginia), Dr. M. Wu (Johns Hopkins University), Dr. M. Zlatic and Dr. C. Eschbach (HHMI Janelia Research Campus), Dr. M. Gallio (Northwestern University), Dr. A. Kopin (Tufts-New England Medical Center), Dr. S. Tanda (Ohio University), Dr. B. Condron (University of Virginia), Dr. T. Kitamoto (University of Iowa) for their kind gift of fly strains.

## Data Availability

Data will be made available on request.

## Supplemental information

**Figure S1.**
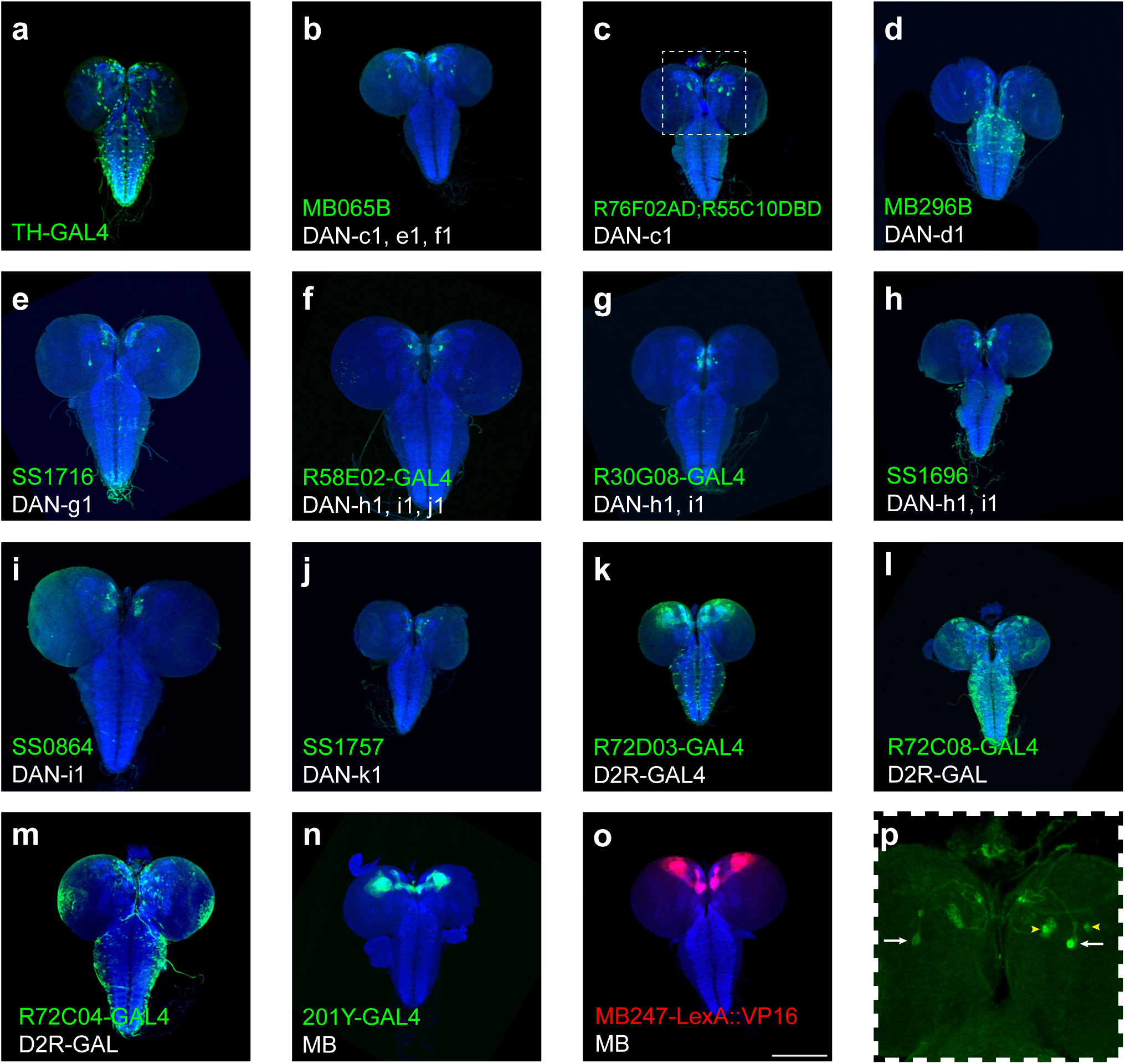
GFP expression patterns in the larval CNS by GAL4 driver strains used in this study. **(a-n)** Representative pictures show the patterns of distinct GAL4 driver strains crossed with a UAS-GFP strain. **(o)** A representative picture shows the pattern of MB247-LexA driver (MB247-LexA::VP16) crossed with a LexAop-RFP strain (LexAop-rCD2::RFP). Blue channel represents neuropils marked by nc82 antibody. Driver strains in **(a-j)** were used in Figure 1d-m. Driver strains in **(k-m)** are strains used in supplementary **Figure S3**. 201Y-Gal4 (**n**) is the driver strain used in Figure 6. Square region is enlarged in **(p)** to show the soma and neurites. White arrows label the soma of DAN-c1, while yellow arrow heads label other neurons. Summary of analysis can be found in Table S3. Scale bar: 200 µm. ***(Note)*** *N numbers can be found in Table S2. In* Figure 1 *and S1, we mainly showed the strains labeling distinct pairs of dopaminergic neurons. The labeling patterns of the rest 47 strains screened were summarized in Table 1, whose brain images are available and can be provided upon request*.

**Figure S2.**
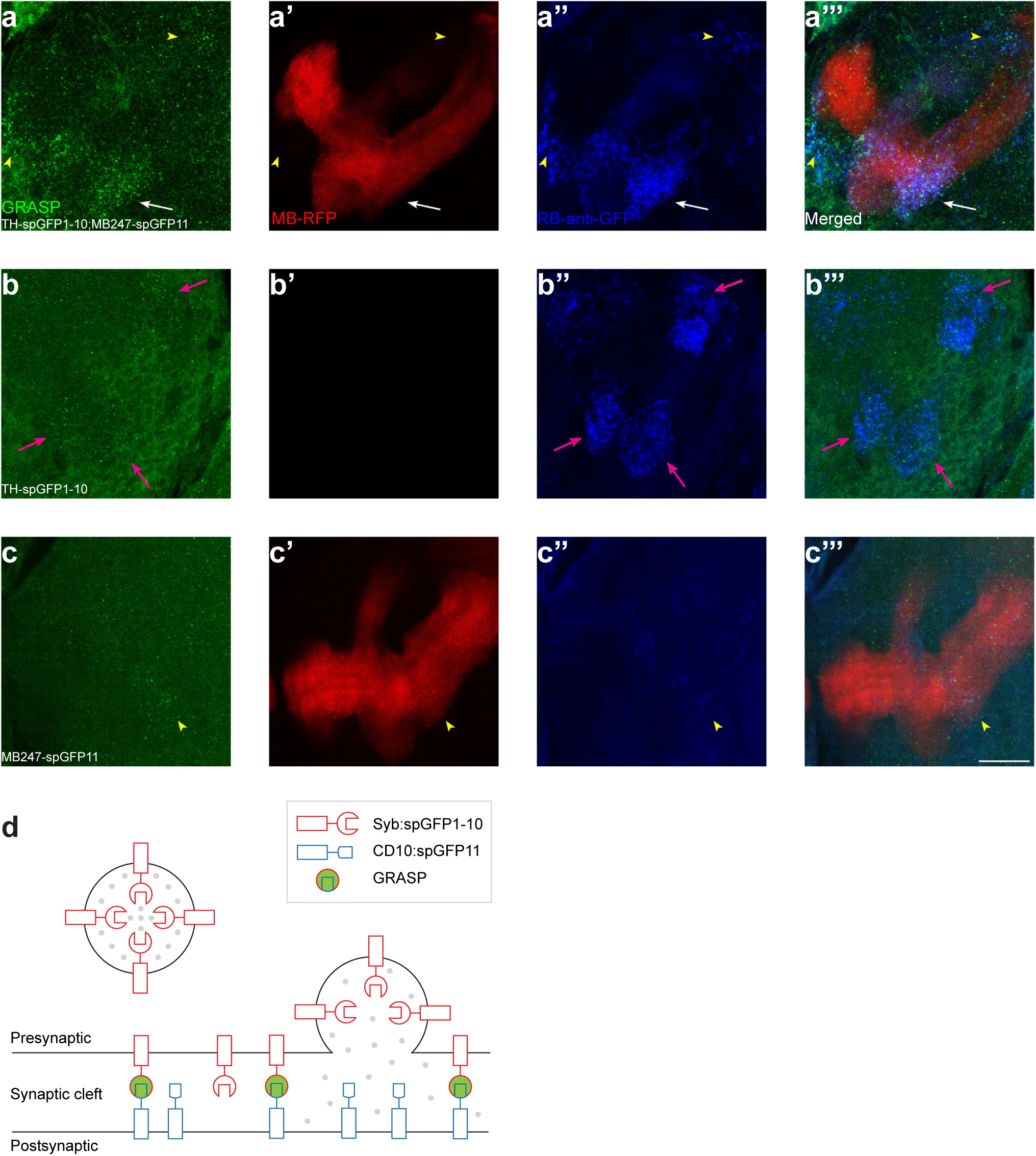
Controls for GRASP experiments. **(a-c)** The GRASP signals were not observed in control groups. The first column shows the mouse anti-GFP staining (green) for GRASP signals, the second column shows MB lobes by RFP (red), and the third column shows the rabbit anti-GFP staining (blue) which only recognizes the spGFP1-10. **(a)** Both GRASP (mouse antibody, green) and spGFP1-10 (rabbit antibody, blue) can be recognized in the lower peduncle (LP) compartment (white arrows), in a larval brain expressing spGFP1-10 under TH-Gal4 and spGFP11 under MB247-LexA. Some strong spGFP1-10 signals outside of MB are also colocalized with GRASP signals (yellow arrowheads). **(b)** GRASP signals are hardly observed in a larval brain only expressing spGFP1-10 under TH-Gal4 (no MB247-LexA in this brain), while anti-spGFP1-10 signals are still strong (magenta arrows). **(c)** Neither GRASP nor spGFP1-10 can be recognized in a larval brain only expressing spGFP11 under MB247-LexA (cyan arrowheads). Scale bars: 20 µm. **(d)** A Schematic diagram shows the mechanism of GRASP. Modified from Macpherson et. al.^50^. *(Note) N numbers can be found in Table S2*.

**Figure S3.**
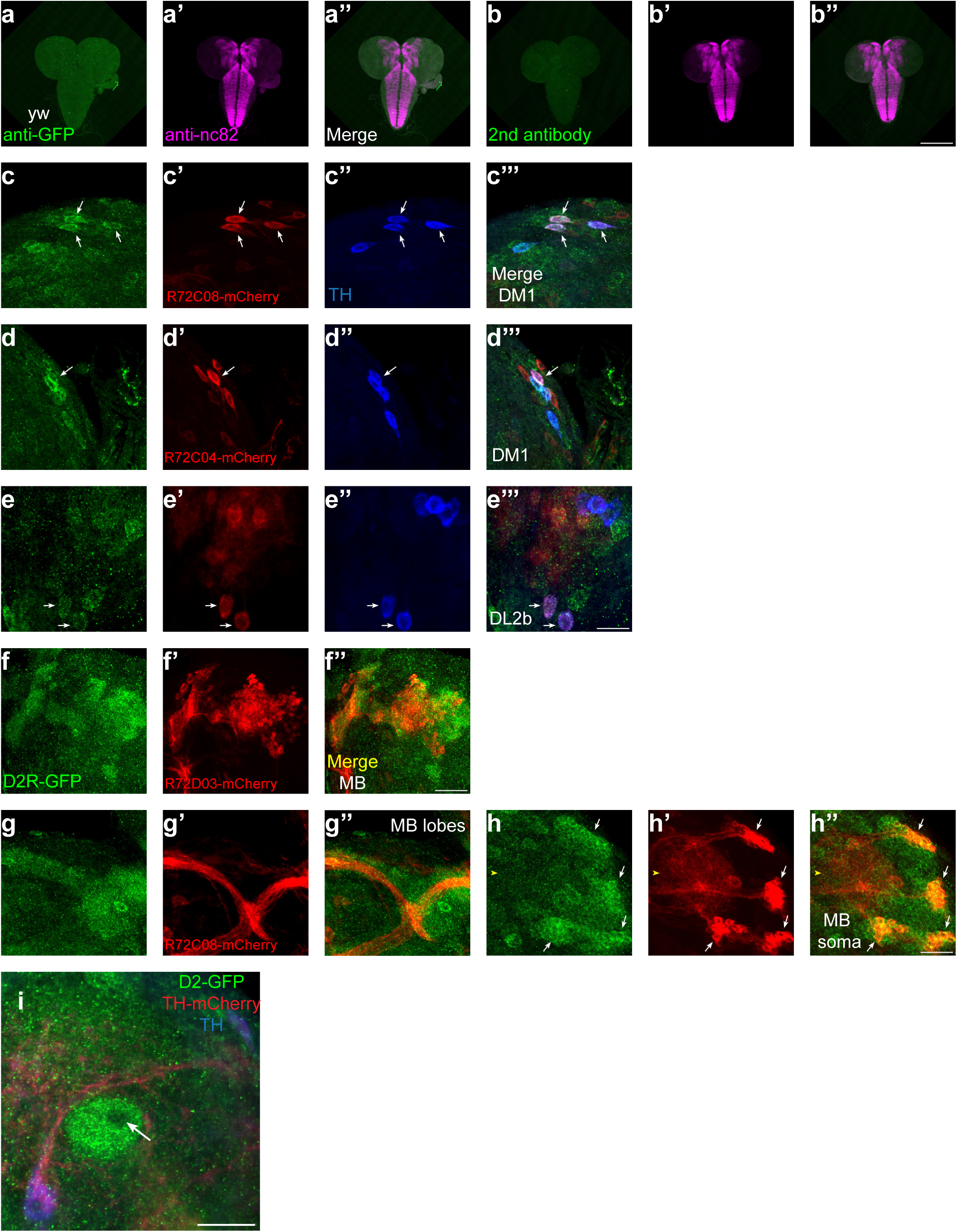
D2R-GAL4 strains support the expression pattern of D2R. **(a-b)** Control staining for D2R-tagged GFP in Figure 3. The green channel represents GFP signals whereas magenta represents neuropils marked by nc82 antibody. GFP staining in WT (a). Secondary antibody only staining for the D2R-tagged GFP strain (b). **(c-e)** D2R-GAL4 strains support the expression of D2R in some DANs. The green channel represents GFP signals, red represents mCherry signals under D2R-GAL4 drivers, and blue marks DANs with TH antibodies. R72C08 labels three DM1 neurons (c). R72C04 labels one DM1 (d) and two DL2b neurons (e). **(f-h)** D2R-GAL4 strains support the expression of D2R in mushroom body neurons. The green channel represents GFP signals, and red represents mCherry signals under D2R-GAL4 drivers. R72D03 marks some MBNs in mushroom body (f). R72C08 labels some MBNs in axons (g), dendrites and soma (h). **(i)** D2R is absent in the core of mushroom body lobes, from a transection view of the lower peduncle. Scale bars: 200 µm (a and b); 20 µm (c-e, g-i), 50 µm (f). *(Note) N numbers can be found in Table S2*.

**Figure S4.**
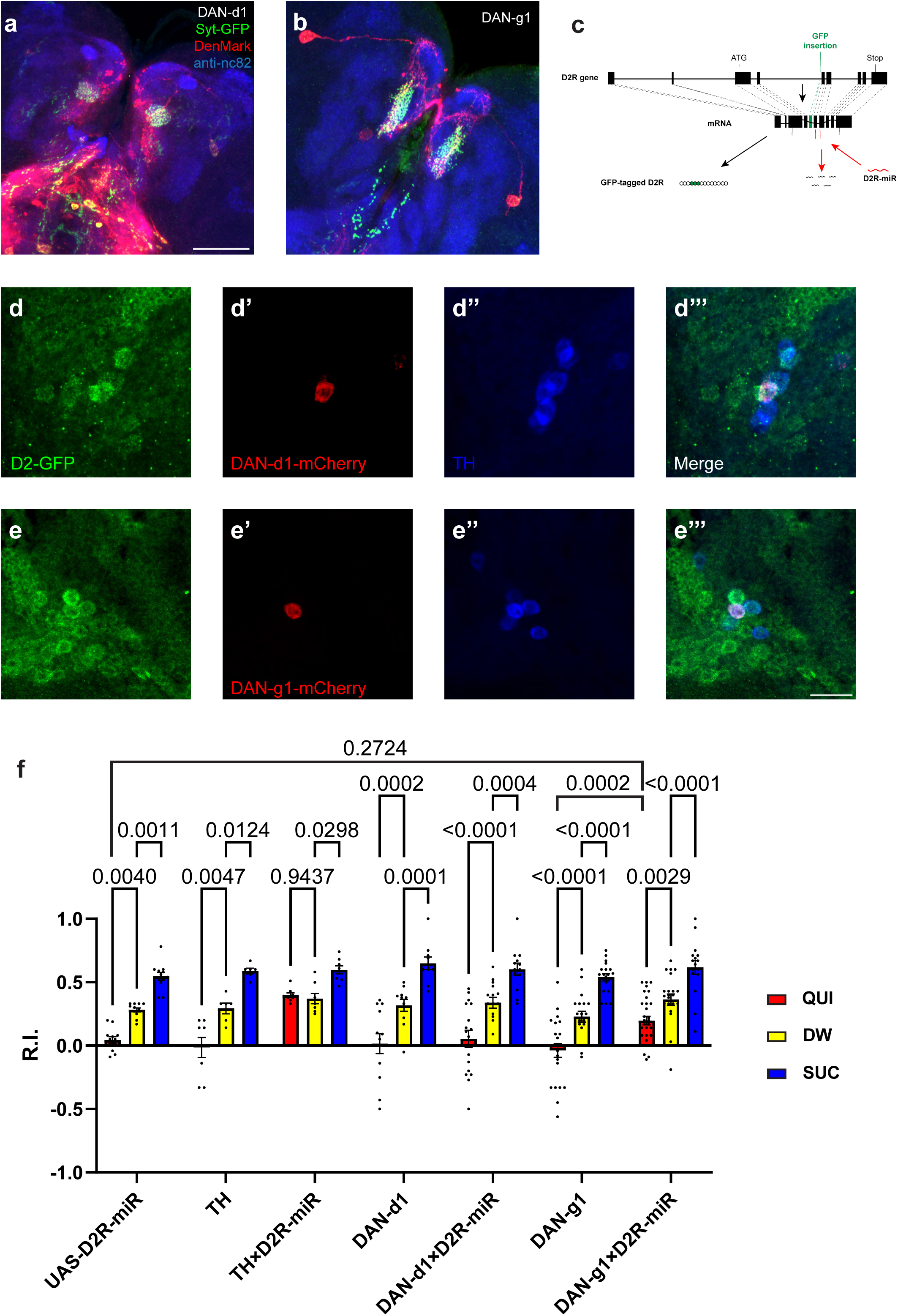
D2Rs in DAN-d1 and DAN-g1 are not necessary for aversive learning. **(a-b)** Dendrites and axons of DAN-d1 (a) and g1 (b) are labeled by DenMark (red) and sytGFP (green), correspondingly. **(c)** The gene structure of D2R, the insertion site of GFP, and the D2R-microRNA targeting sequence are shown in a schematic diagram. Modified from Xie and Ho^60^. **(d-e)** D2Rs are expressed in DAN-d1 (d) and DAN-g1 (e). The expression pattern of D2R is shown with tagged-GFP, DAN-g1 and d1 are marked by mCherry, and DANs are labeled by TH-antibody (blue). **(f)** Knockdown D2R in neither in DAN-g1 nor DAN-d1 impairs aversive learning. Data are shown as mean ± SEM. UAS-D2R-miR data is from Figure 4e. Two-way ANOVA, Dunnett’s multiple comparison test. p = 0.0544 for interaction, p < 0.0001 for row factor (genotype), and p < 0.0001 for column factor (US). For N numbers, see Table S5. Scale bars: 50 µm (a and b), 20 µm (d and e). *(Note) N numbers of immunostaining for each strain can be found in Table S2*.

**Figure S5.**
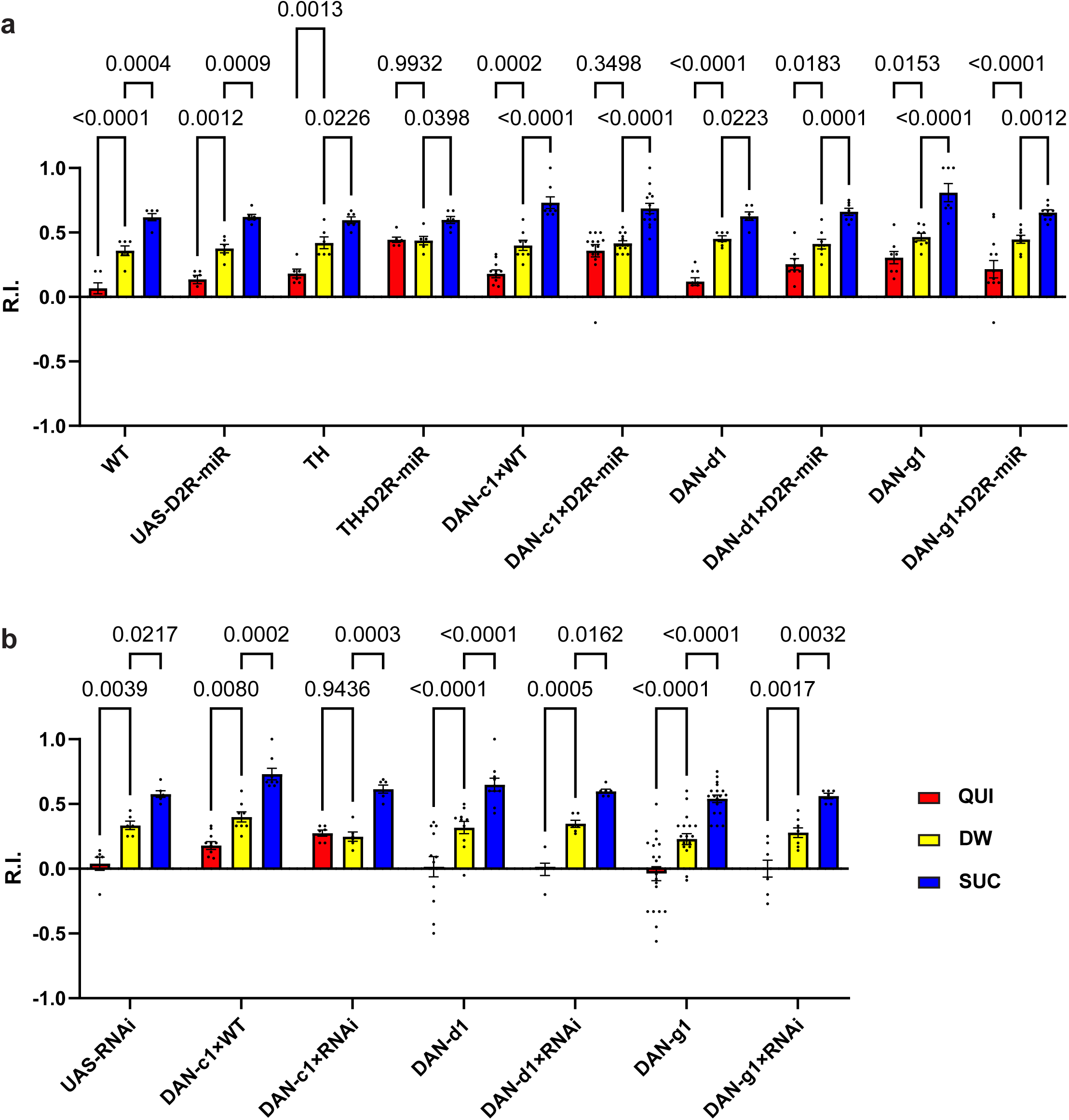
Knockdown D2R in DAN-c1 with a RNAi strain also impairs aversive learning. **(a)** Knockdown of D2R with D2R-miR in DAN-c1 impairs larval aversive learning when using propionic acid as the odorant. **(b)** Knockdown of D2R in DAN-c1 with a RNAi strain also impairs larval aversive learning when using pentyl acetate as the odorant, while knockdown of D2R in either DAN-d1 or g1 does not affect learning. Data are shown as mean ± SEM. Two-way ANOVA, Dunnett’s multiple comparison test. In D2R-miR experiments (a), p = 0.0009 for interaction, p < 0.0001 for row factor (genotype), and p < 0.0001 for column factor (US); in D2R-RNAi experiments (b), p = 0.3802 for interaction, p = 0.0001 for row factor (genotype), and p < 0.0001 for column factor (US). For N numbers, see Table S5.

**Figure S6.**
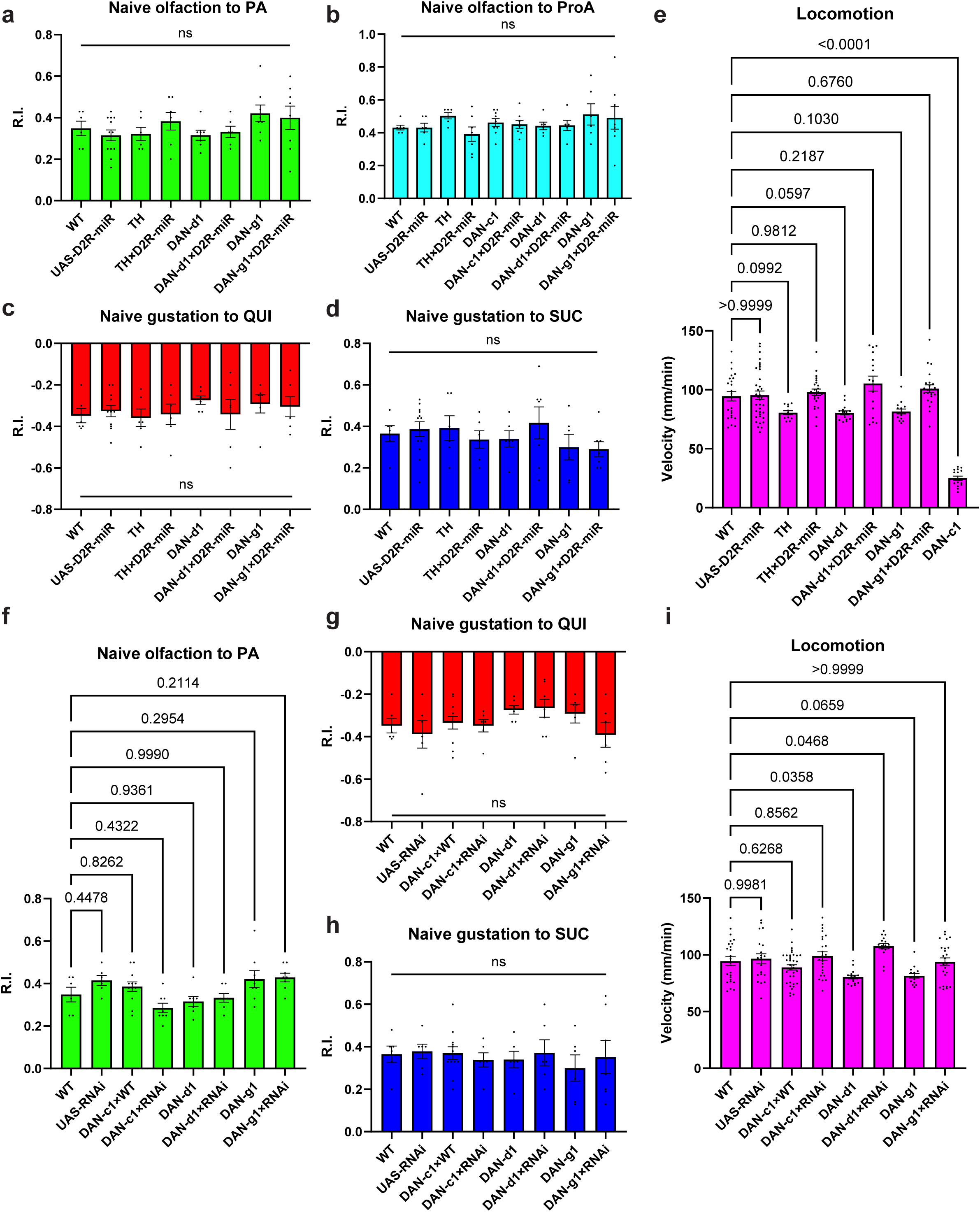
Naïve sensory and motor functions of *Drosophila* larvae. **(a-e)** The naïve sensory and motor functions in larvae related to D2R-miR experiments. **(a)** No significant difference exists between strains in naïve olfactory tests toward pentyl acetate (PA). **(b)** No significant difference exists between strains in naïve olfactory tests toward propionic acid (ProA). **(c)** No significant difference exists between strains in naïve gustatory tests toward quinine (QUI). **(d)** No significant difference exists between strains in naïve gustatory tests toward sucrose (SUC). **(e)** No significant difference is found between strains and WT, except DAN-c1. So, larvae from DAN-c1 × WT are used for all behavioral assays (refer to **i**). **(f-i)** The naïve sensory and motor functions in larvae related to RNAi experiments. **(f)** No significant difference exists between strains and WT. **(g)** No significant difference exists between strains in naïve gustatory tests toward QUI. **(h)** No significant difference exists between strains in naïve gustatory tests toward SUC. **(i)** There is a significant difference between strains in larval locomotion speed, and there is significant difference between DAN-d1, DAN-d1 × RNAi and WT, but these differences do not affect the learning ability of larvae. Data are shown as mean ± SEM. One-way ANOVA, Dunnett’s multiple comparison test. For N numbers, see Table S5.

**Figure S7.**
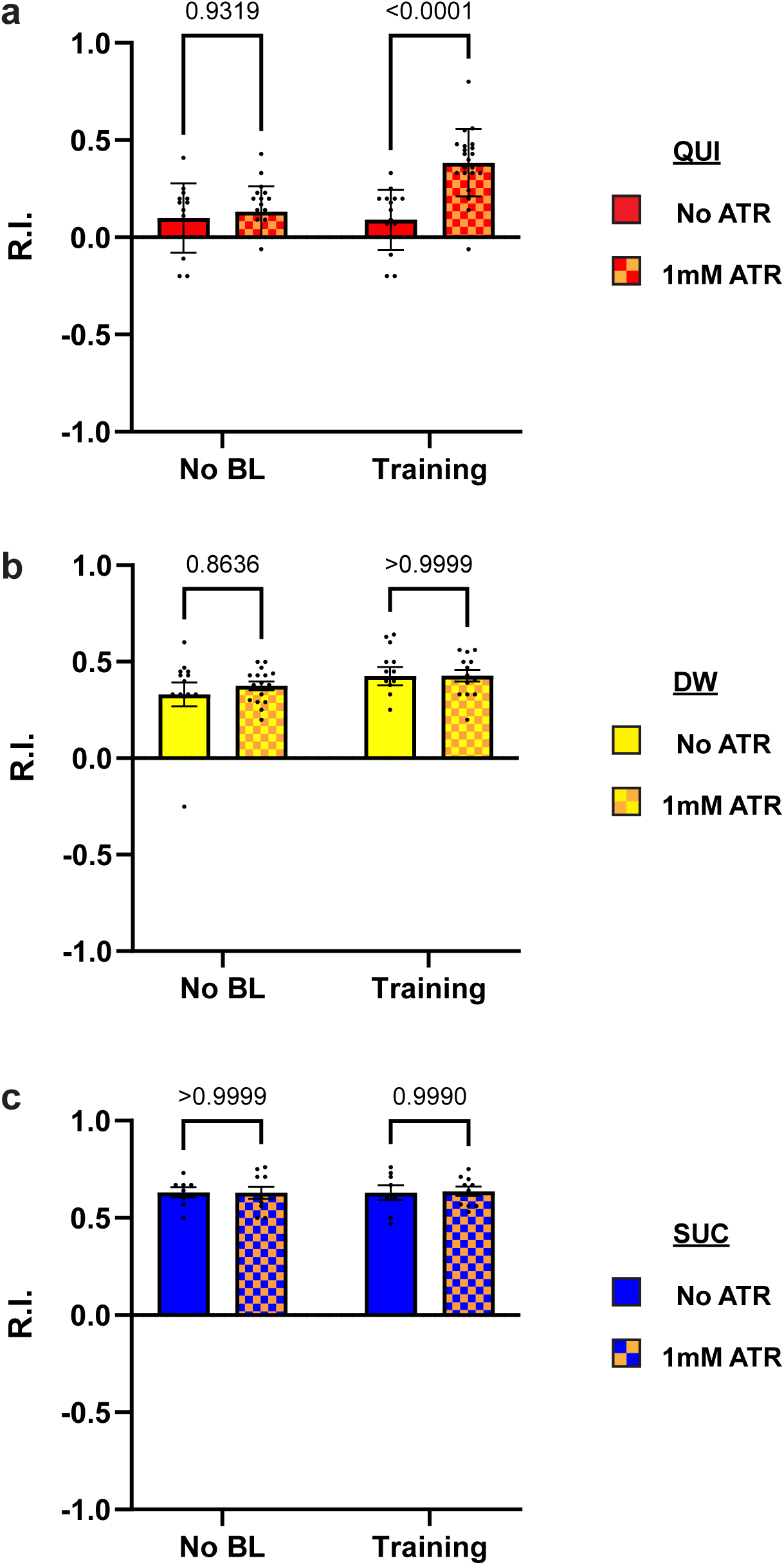
Over-excitation of DANs during learning impairs larval aversive learning. Third-instar larvae with ChR2 expression in most DANs (TH-Gal4) were used. Activation of DANs during training impairs larval aversive learning (**a**), while keeping appetitive learning intact (**c**). No learning behaviors are observed in the control distilled water (DW) groups **(b)**. Data are shown as mean ± SEM. Two-way ANOVA, Tukey’s multiple comparison test. In QUI group (a), p = 0.0011 for interaction, p = 0.0023 for row factor (training stages), and p < 0.0001 for column factor (whether with ATR); in DW group (b), p = 0.6126 for interaction, p = 0.0850 for row factor (training stages), and p = 0.5748 for column factor (whether with ATR); in SUC group (c), p = 0.8910 for interaction, p = 0.9239 for row factor (training stages), and p = 0.9503 for column factor (whether with ATR). For N numbers, see Table S6.

### Supplementary Tables

**Table S1.** Other strains used in this study.

|  | Strains | Information | Source/Gift | Stock # |
| --- | --- | --- | --- | --- |
| 1 | R72C04 | D2R-GAL4 driver (verified) | BDSC | 49416 |
| 2 | R72C08 | D2R-GAL4 driver (verified) | BDSC | 49621 |
| 3 | R72D03 | D2R-GAL4 driver | BDSC | 46676 |
| 4 | D2R-EGFP | EGFP-tagged D2R | BDSC | 60276 |
| 5 | 201Y-GFP | GFP expressed in Mushroom Body | BDSC | 64296 |
| 6 | MB247-LexA::Vp16/TM6B | MB-LexA, mushroom body driver | Dr. M. Gallio's Lab | Macpherson et al. 2015 |
| 7 | UAS-nSyb-spGFP1-10, LexAop-CD4-spGFP11/CyO | GRASP strain | BDSC | 64314 |
| 8 | LexAop-nSyb-spGFP1-10, UAS-CD4-spGFP11; MKRS/TM6B | Reverse GRASP strain | BDSC | 64315 |
| 9 | WT(Canton-S) | Wild type | BDSC | 64349 (1) |
| 10 | UAS-RNAi(D2R) (III) line 2 | UAS-D2R-RNAi | Dr. A. Kopin's Lab | Draper et al. 2007 |
| 11 | 201Y-GAL4 | MB-GAL4, mushroom body driver | BDSC | 4440 |
| 12 | CyO/Sco; TM2, Ubx/TM6, Hu, Tb | Ballancer Chromosome | Dr. S. Tanda's Lab |  |
| 13 | 1407-Gal4 | Pan-neuronal driver | BDSC | 8751 |
| 14 | 10XUAS-mCD8::GFP (I) | GFP strain | BDSC | 32189 |
| 15 | 10XUAS-mCD8::GFP (II) | GFP strain | BDSC | 32186 |
| 16 | 10XUAS-mCD8::GFP (III) | GFP strain | BDSC | 32184 |
| 17 | UAS-Chr2; UAS-ChrR2 | Optogenetics strain | Dr. B. Condron's Lab |  |
| 18 | LexAop-rCD2::RFP; UAS-CD4-spGFP1-10, LexAop-CD4-spGFP11 | RFP expressed in MB; GRASP | BDSC | 58755 |
| 19 | UAS-CD8::mCherry (II) | mCherry strain | BDSC | 27391 |
| 20 | UAS-CD8::mCherry (III) | mCherry strain | BDSC | 27392 |
| 21 | yw1118 | Wild type for D2R-EGFP | Dr. S. Tanda's Lab |  |
| 22 | D2R-miR | UAS-DD2R-microRNA | Dr. M. Wu's Lab | Xie et al. 2018 |
| 23 | UAS-*Shibirets1* | Thermogenetics strain | Dr. T. Kitamoto's Lab |  |
| 24 | UAS-dTRPA1 | Thermogenetics strain | BDSC | 26263 |
| 25 | UAS-nsyb::GFP; UAS-DenMark | Pre- and postsynaptic terminals | BDSC | 33056 |

**Table S2.** N numbers for brain samples used in Figures.

| **Figure 1** | **N** |  |  | **Figure S1** | **N** |
| --- | --- | --- | --- | --- | --- |
| **Strains** | **IHC** | **GRASP** |  | TH-GAL4 | 4 |
| TH-GAL4 | 8 | 16 |  | MB065B | 4 |
| MB065B | 14 | 15 |  | R76F02AD;R55C10DBD | 5 |
| R76F02AD;R55C10DBD | 13 | 13 |  | MB296B | 3 |
| MB296B | 14 | 4 |  | SS1716 | 2 |
| SS1716 | 10 | 21 |  | R58E02 | 3 |
| R58E02 | 5 | 10 |  | R30G08 | 2 |
| R30G08 | 13 | 6 |  | SS1696 | 4 |
| SS1696 | 16 | 4 |  | SS0864 | 4 |
| SS0864 | 7 | 8 |  | SS1757 | 4 |
| SS1757 | 6 | 13 |  | R72D03-GAL4 | 7 |
|  |  |  |  | R72C08-GAL4 | 5 |
| **Figure 2** | **N** |  |  | R72C04-GAL4 | 3 |
| DAN-c1 x Syt-GFP;DenMark | 4 |  |  | 201Y-GAL4 | 4 |
|  |  |  |  | MB247-LexA::VP16 | 3 |
| **Figure 3** | **N** |  |  |  |  |
| DM1a | 11 |  |  | **Figure S2** | **N** |
| DM1b | 9 |  |  | TH-spGFP1-10 | 3 |
| pPAM | 11 |  |  | MB247-spGFP11 | 4 |
| DL1 | 10 |  |  |  |  |
| DL2a | 7 |  |  | **Figure S3** | **N** |
| DL2b | 5 |  |  | R72D03 | 5 |
| MB Soma | 6 |  |  | R72C08 | 4 |
| MB Lobe | 8 |  |  | R72C04 | 5 |
| **Figure 4** | **N** |  |  | **Figure S4** | **N** |
| DAN-c1 x D2-GFP | 9 |  |  | DAN-d1 x Syt-GFP;DenMark | 6 |
| TH-Gal4 | 7 |  |  | DAN-g1 x Syt-GFP;DenMark | 3 |
| TH-miR | 6 |  |  | DAN-d1 x D2-GFP | 13 |
|  |  |  |  | DAN-g1 x D2-GFP | 7 |

**Table S3.** Summary of R76F02AD;R55C10DBD (DAN-c1) identifying patterns in larval brains. For the strain R76F02AD; R55C10DBD, 22 third-instar larval brains expressing GFP or SytGFP and DenMark were examined, and all of them clearly identified DAN-c1. Half of them only identified DAN-c1, the rest had 1 to 5 weak identified cells without neurites, and barely 1 or 2 strong identified cells appeared. These non-DAN-c1 neurons were seldom dopaminergic neurons. In ventral nerve cord (VNC), 8 out of 12 did not have any identified cells, 3 had 2-4 strong identified cells. These data supported that R76F02AD;R55C10DBD exclusively labels DAN-c1 in third-instar larval brains.

| **R76F02AD;R55C10DBD (DAN-c1) brains** | **other DAN** | **None DAN** | **VNC** |
| --- | --- | --- | --- |
| **1** | 0 | 1 strong, 4 weak | NA |
| **2** | 1 weak no neurites | 0 | 3 weak |
| **3** | 0 | 7 weak | NA |
| **4** | 0 | 0 | 2 strong |
| **5** | 0 | 0 | 2 strong |
| **6** | 0 | 2 strong, 3 weak | NA |
| **7** | 0 | 0 | NA |
| **8** | 0 | 0 | 4 strong |
| **9** | 0 | 5 weak | NA |
| **10** | 0 | 1 weak | NA |
| **11** | 0 | 8 weak | NA |
| **12** | 0 | 0 | NA |
| **13** | 0 | 4 strong 1 weak | NA |
| **14** | 0 | 0 | 0 |
| **15** | 0 | 0 | 0 |
| **16** | 0 | 0 | NA |
| **17** | 0 | 0 | 0 |
| **18** | NA | 1 strong, 1 weak | 0 |
| **19** | NA | 7 weak | 0 |
| **20** | NA | 5 weak | 0 |
| **21** | NA | 0 | 0 |
| **22** | NA | 1 weak | 0 |

**Table S4.** N numbers and p-values for learning assays with thermogenetics.

|  | 22°C |  |  | 34°C |  |  | Interaction<br>p-value | Row Factor p-value<br>(Temperature) | Column<br>Factor<br>p-value<br>(US) |
| --- | --- | --- | --- | --- | --- | --- | --- | --- | --- |
|  | QUI | DW | SUC | QUI | DW | SUC |  |  |  |
| **shibire<sup>ts1</sup>** | 9 | 8 | 8 | 9 | 9 | 9 | 0.8880 | 0.5290 | <0.0001 |
| **DAN-**<br>**c1×shibire<sup>ts1</sup>** | 14 | 6 | 7 | 17 | 6 | 9 | <0.0001 | 0.0018 | <0.0001 |
| **DAN-c1** | 8 | 9 | 9 | 9 | 9 | 9 | 0.6143 | 0.1664 | <0.0001 |
| **DAN-**<br>**c1×dTRPA1** | 15 | 14 | 13 | 14 | 15 | 13 | <0.0001 | 0.0019 | <0.0001 |
| **dTRPA1** | 9 | 8 | 9 | 10 | 10 | 10 | 0.8001 | 0.4228 | <0.0001 |

**Table S5.**
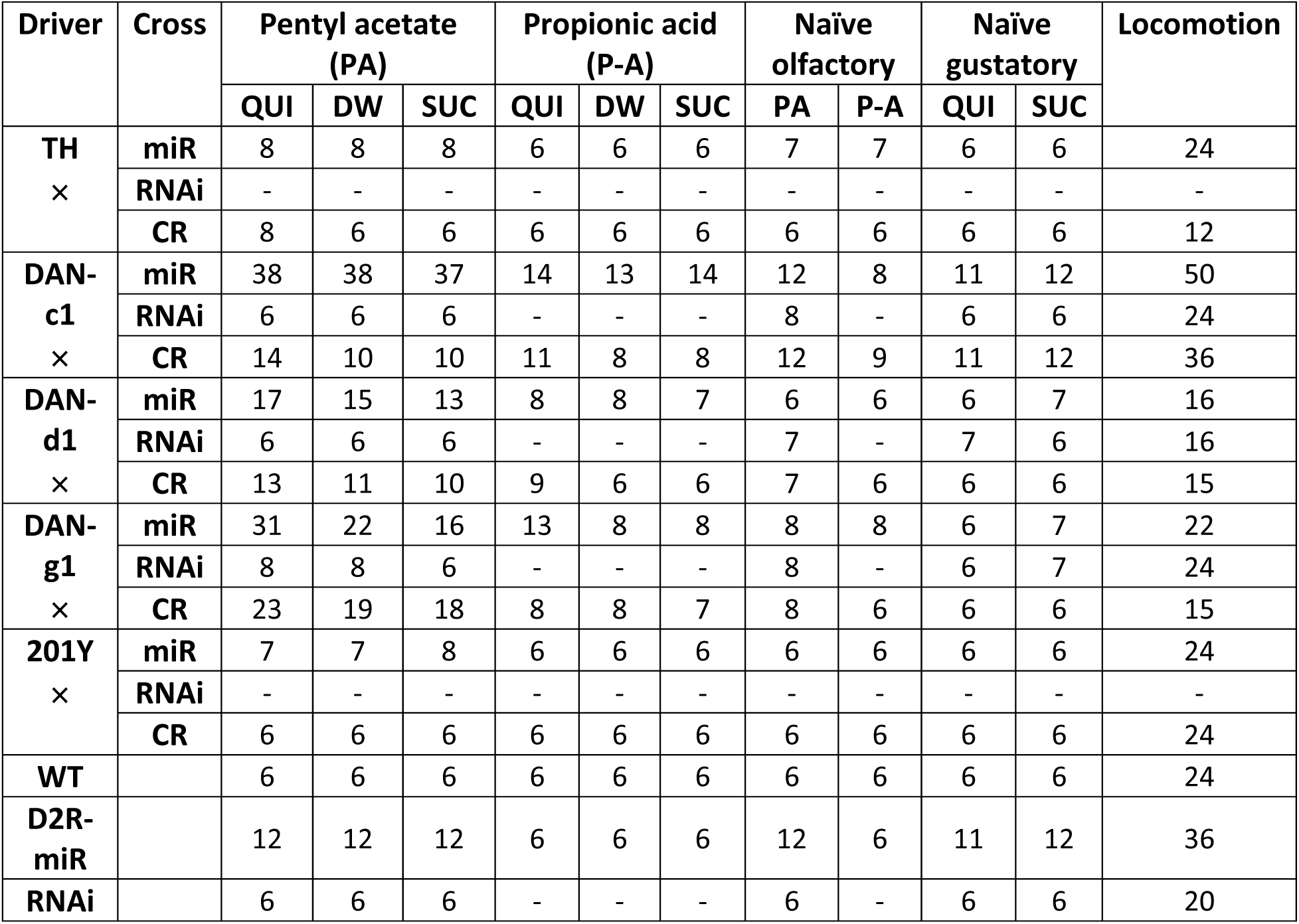
N numbers for D2R knockdown experiments.

**Table S6.** N numbers for learning assays with optogenetics.

| Strains | Groups | BL |  |  |  |  |  |  |  |  | No BL |  |  |
| --- | --- | --- | --- | --- | --- | --- | --- | --- | --- | --- | --- | --- | --- |
|  |  | Resting |  |  | Learning |  |  | Testing |  |  |  |  |  |
|  |  | QUI | DW | SUC | QUI | DW | SUC | QUI | DW | SUC | QUI | DW | SUC |
| TH×Chr2 | ATR | - | - | - | 21 | 13 | 10 | - | - | - | 20 | 16 | 10 |
|  | No ATR | - | - | - | 16 | 13 | 8 | - | - | - | 14 | 13 | 8 |
| DAN-c1×Chr2 | ATR | 11 | 8 | 9 | 15 | 12 | 9 | 9 | 8 | 8 | 16 | 11 | 9 |
|  | No ATR | 9 | 9 | 8 | 15 | 12 | 8 | 9 | 9 | 9 | 16 | 12 | 9 |
| 201Y×Chr2 | ATR | 8 | 7 | 7 | 9 | 6 | 9 | 7 | 7 | 10 | 11 | 8 | 7 |
|  | No ATR | 10 | 7 | 7 | 10 | 7 | 7 | 9 | 7 | 13 | 12 | 10 | 7 |

